# HMMploidy: inference of ploidy levels from short-read sequencing data

**DOI:** 10.1101/2021.06.29.450340

**Authors:** Samuele Soraggi, Johanna Rhodes, Isin Altinkaya, Oliver Tarrant, François Balloux, Matthew C. Fisher, Matteo Fumagalli

## Abstract

The inference of ploidy levels from genomic data is important to understand molecular mechanisms underpinning genome evolution. However, current methods based on allele frequency and sequencing depth variation do not have power to infer ploidy levels at low-and mid-depth sequencing data, as they do not account for data uncertainty. Here we introduce `HMMploidy`, a novel tool that leverages the information from multiple samples and combines the information from sequencing depth and genotype likelihoods. We demonstrate that `HMMploidy` outperforms existing methods in most tested scenarios, especially at low-depth with large sample size. We apply `HMMploidy` to sequencing data from the pathogenic fungus *Cryptococcus neoformans* and retrieve pervasive patterns of aneuploidy, even when artificially downsampling the sequencing data. We envisage that `HMMploidy` will have wide applicability to low-depth sequencing data from polyploid and aneuploid species.

## Introduction

In recent years, advances in Next Generation Sequencing (NGS) technologies allowed for the generation of large amount of genomic data (Levy and Myers, 2016; Metzker, 2010). Many statistical and computational methods, and accompanying software, to process NGS data for genotype and variant calling have been proposed (Garrison and Marth, 2012; Li et al., 2009; Van der Auwera et al., 2013). Additionally, dedicated software have been developed to analyse low-coverage sequencing data (Fumagalli, Vieira, Linderoth, et al., 2014; Nielsen et al., 2011), a popular and cost-effective approach in population genomic studies (Lou et al., 2021). However, most of these efforts have been focused towards model species with known genomic information. In particular, there has been a lack of research into modelling sequencing data from non-diploid species or organisms with unknown ploidy.

Polyploidy is typically defined as the phenomenon whereby the chromosome set is multiplied, resulting the organism to have three or more sets of chromosomes (Van de Peer et al., 2017). Polyploidy is common to many organisms, and it can be the consequence of hybridisation or whole genome duplication (Fox et al., 2020). For instance, polyploidy plays a significant role in the evolution and speciation of plants (Sattler et al., 2016), as 34.5% of vascular plants (including leading commercial crop species) are shown to be polyploid (Wood et al., 2009).

Of particular interest is the case of aneuploidy, whereby chromosomal aberrations cause the number of chromosomal copies to vary within populations and individuals. Ploidy variation can be associated with a response or adaptation to environmental factors (Coward and Harding, 2014), and it is a phenomenon commonly detected in cancer cells (Davoli and Lange, 2011) and several pathogenic fungi (i.e. *Cryptococcus neoformans, Candida albicans* and *Candida glabrata*) and monocellular parasites (Avramovska et al., 2021; Farrer et al., 2013; Fu et al., 2021; Morrow and Fraser, 2013; Stone et al., 2019; Yang et al., 2021; Zhu et al., 2018).

Among aneuploid species, *Cryptococcus neoformans* is a fungal pathogen capable of causing meningitis in immunocompromised individuals, particularly HIV/AIDS patients (May et al., 2016). Ploidy variation, via aneuploidy and polyploidy, is an adaptive mechanism in *Cryptococcus neoformans* capable of generating variation within the host in response to a harsh environment and drug pressure (Morrow and Fraser, 2013). Aneuploidy-driven heteroresistance to the frontline antifungal drug fluconazole has been described (Stone et al., 2019), resulting in treatment failure in patients. Within fluconazole resistant colonies, aneuploidy was common, particularly disomy of chromosome 1 which harbours the gene encoding the main drug target of fluconazole, *ERG11* (Stone et al., 2019). For these reasons, inferring the ploidy of a sample from genomic data, like in the case of *Cryptococcus neoformans*, is essential to shed light onto the evolution and adaptation across the domains of life.

Available computational methods to infer ploidy levels from genomic data are based either on modelling the distribution of observed allele frequencies (nQuire (Weiß et al., 2018)), comparing frequencies and coverage to a reference data set (`ploidyNGS` (Augusto Corrêa dos Santos et al., 2017)), or using inferred genotypes and information on GC-content, although the latter is an approach specific for detecting aberrations in cancer genomes (e.g. `AbsCN-seq` (Bao et al., 2014), sequenza (Favero et al., 2015)). A popular approach is based on the simple eyeballing method, that is, on the visual inspection of variation of sequencing depth (compared to another ground-truth data set sequenced with the same setup) and allele frequencies (Augusto Corrêa dos Santos et al., 2017). However, methods based only on sequencing depth, allele frequencies and genotypes limit the inference on the multiplicity factor of different ploidy levels only (if present). Additionally, they often need a reference data with known ploidy to be compared to, and they generally lack power for low-or mid-depth sequencing data applications, which are typically affected by large data uncertainty. As low-coverage whole genome sequencing is a common strategy in population genetic studies of both model and non-model species (Therkildsen and Palumbi, 2017), a tool that incorporates data uncertainty is in dire need.

To overcome these issues, we introduce a new method called `HMMploidy` to infer ploidy levels from low-and mid-depth sequencing data. `HMMploidy` comprises a Hidden Markov Model (HMM) (Rabiner, 1989) where the emissions are both sequencing depth levels and observed reads. The latter are translated into genotype likelihoods (Nielsen et al., 2011) and population frequencies to leverage the genotype uncertainty. The hidden states of the HMM represent the ploidy levels which are inferred in windows of polymorphisms. Notably, `HMMploidy` determines automatically its number of latent states through a heuristic procedure and reduction of the transition matrix. Moreover, our method can leverage the information from multiple samples in the same population by estimate of population frequencies, making it effective at very low depth.

`HMMploidy` infers ploidy variation in sliding windows among chromosomes and among individuals. While ploidy is not expected to vary within each chromosome, the distribution of inferred ploidy tracts provides further statistical support to whole-chromosome estimates. Additionally, `HMMploidy` can identify local regions with aberrant predicted ploidy to be further investigated, for instance as potential locations of copy number variants (CNVs) or structural rearrangements. Finally, any detected within-chromosome ploidy variation can serve as a diagnostic tool to investigate possible mapping or assembly errors. Notably, by training separate HMMs, `HMMploidy` can effectively infer aneuploidy among chromosomes and samples.

`HMMploidy` is written in `R/C++` and `python`. Source code is freely available at https://github.com/SamueleSoraggi/HMMploidy, integrated into ngsTools (Fumagalli, Vieira, Linderoth, et al., 2014), and FAIR data sharing is available at the OSF repository https://osf.io/5f7ar. We will first introduce the mathematical and inferential model underlying `HMMploidy`, then show its performance to detect ploidy levels compared to existing tools, and finally illustrate an application to sequencing data from the pathogenic fungus *Cryptococcus neoformans*.

## Material and methods

This section describes the methods used in the implementation of the `HMMploidy` software. In what follows, data is assumed to be diallelic (i.e. we observe at most two states at a particular genotype regardless of the number of copies), without loss of generality. Allowing for more than two alleles would add a summation over all possible pairs of alleles in all calculations. In our notation, indices are lower case and vary within an interval ranging from 1 to the index’s upper case letter, e.g. *m* = 1, …, *M*.

### Probability of sequenced data

Let *O* = (*O*_1_, …, *O*_*M*_) be the observed NGS data for *M* sequenced genomes at *N* sites. Consider an *m*-th genome and *n*-th locus. We define a locus as a nucleotide site. We assume that sequencing reads are mapped and aligned so that bases can be assigned to a single nucleotide site. For ease of notation, we suppress the two indices, since they do not vary in the formula (1). For such genome and locus define *Y, G* and *O* as the ploidy, genotype and sequencing data, respectively. Given *Y*, the genotype *G* assumes values in {0, …, *Y*}, where each value is the number of alternate (or derived) alleles of said genotype.

The probability of the sequenced data, conditionally on the ploidy *Y* and the population allele frequency *F* at locus *n*, is expressed by

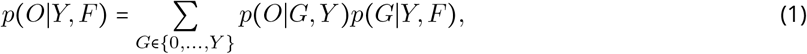

where the left-hand side of the equation has been marginalised over the genotypes, and the resulting probabilities have been rewritten as product of two terms using the tower property of the probability. The first factor of the product is the genotype likelihood (McKenna et al., 2010). Note that the only varying parameter in it is the genotype; therefore it is usually rewritten as *L* (∣*O, Y*). The second factor is the probability of the genotype given the population allele frequency and the ploidy level, in other words the prior probability of the genotype. The marginalisation over all possible genotypes has therefore introduced a factor that takes into account the genotype uncertainty. The calculation of genotype likelihoods for an arbitrary ploidy number and the estimation of population allele frequencies are described in the Supplementary Material.

Throughout the analyses carried out in this paper, we assume Hardy-Weinberg equilibrium (HWE) and thus model the genotype probability with a binomial distribution (Hardy, 1908; Weinberg, 1908). Other methods considering departure from HWE (DHW), can be considered and implemented by *ad hoc* substitutions of the formula coded in the software. Such functions can be useful in specific situations, such as pathology-, admixture-and selection-induced DHW scenarios (Chen et al., 2017; Lachance, 2009; Wittke-Thompson et al., 2005). However, we will leave the treatment of DHW for the inference of ploidy variation to future studies.

### Hidden Markov Model for ploidy inference

Here, the HMM is defined, and the inferential process of ploidy levels from the HMM is illustrated. Further mathematical details, proofs and algorithms are available in the Supplementary Material. Consider the *N* sites arranged in *K* adjacent and non-overlapping windows. For each individual *m*, `HMMploidy` defines a HMM with a Markov chain of length *K* of latent states 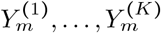, as shown for a sequence of two ploidy levels (Fig. 1A) in the graphical model of dependencies of Fig. 1B. Each *k*-th latent state represents the ploidy level at a specific window of loci, and each window’s ploidy level depends only on the previous one. Therefore, the sequence of states is described by a transition matrix ***A*** of size ∣𝒴∣ × ∣𝒴∣ and a ∣𝒴∣ -long vector of starting probabilities ***δ***, where 𝒴 is the set of ploidy levels included in the model and ∣𝒴∣ is the number of ploidy levels (i.e. cardinality of 𝒴) (Fig. 1C).

**Figure 1. :**
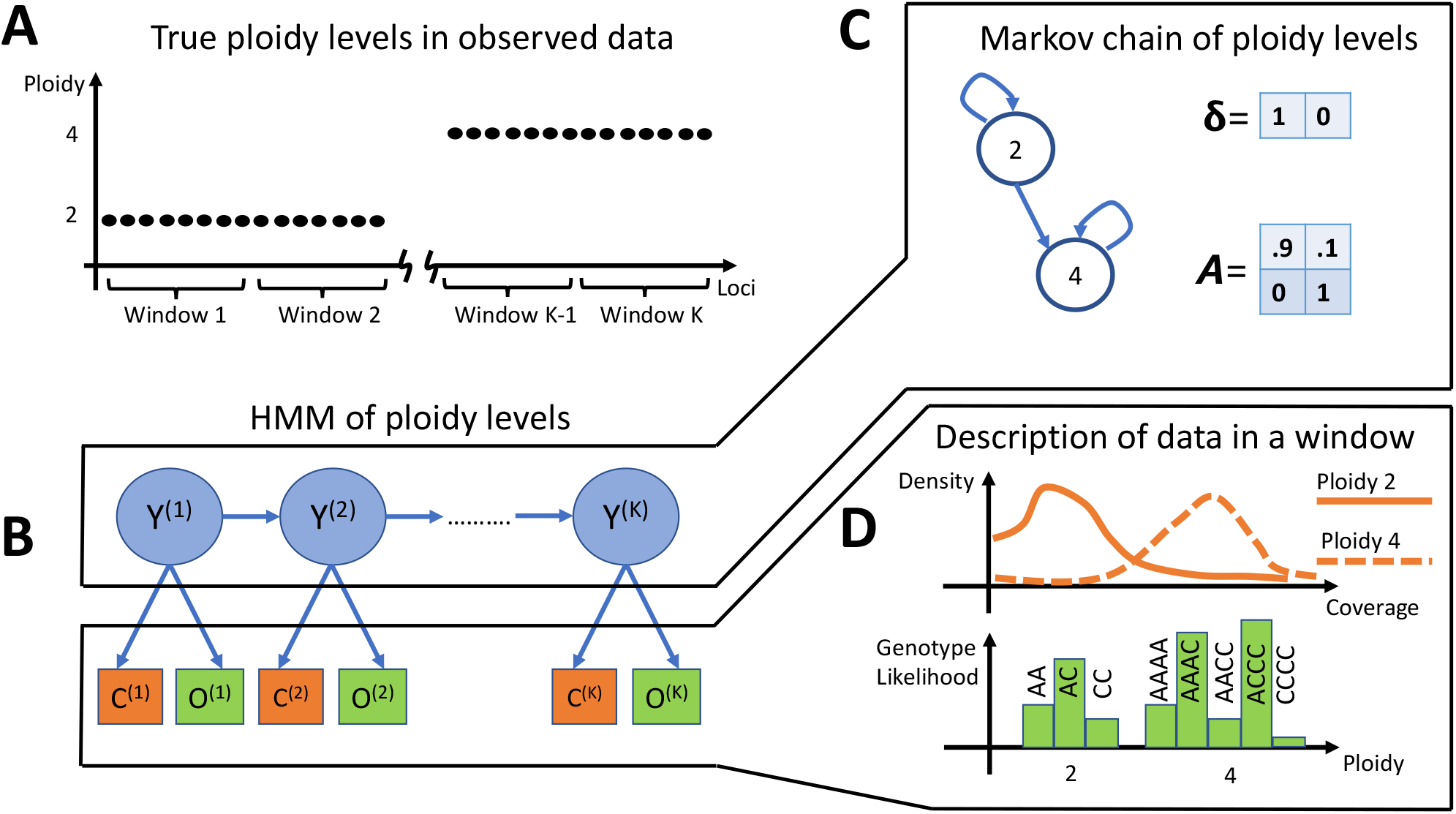
HMM for two ploidy levels. (A) Consider a NGS dataset consisting of a sequence of two ploidy levels. (B) The HMM describing the data has a sequence of hidden states *Y* ^(1)^, …, *Y* ^(*k*)^ - one for each window of loci - that can assume one of two values of the ploidies. Observations *C* ^(1)^, …, *C* ^(*k*)^ and *O* ^(1)^, …, *O* ^(*k*)^ describe the sequencing depth and observed reads in each window, respectively. The index related to the sample is omitted to simplify the notation. (C) The sequence of ploidy levels is described by a Markov chain with two states, governed by a starting vector ***δ*** and a Markov matrix ***A***. (D) At each window, the observations are described by the distribution of depth. There are two distributions, each one dependant on the ploidy level. Similarly, genotype likelihoods describe the observed reads by modelling the genotypes at two distinct ploidy levels.

In the HMM structure, each of the ∣𝒴∣ ploidy levels emits two observations (Fig. S1). Those contain a dependency on which ploidy is assigned to that window. The observations consist of the sequenced reads 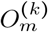 and the average sequencing depth 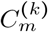 in the *k*-th window (Fig. 1B). The former is modelled by the probability in Equation 1; the latter by a Poisson-Gamma distribution (Bishop, 2006; Casella and Berger, 2002) (Fig. 1D). The Poisson-Gamma distribution consists of a Poisson distribution whose mean parameter is described by a Gamma random variable. This generates a so-called super-Poissonian distribution, for which the mean is lower than the variance. This allows us to model overdispersed counts, a common issue in NGS datasets (Anders and Huber, 2010).

For the *m*-th HMM, the Poisson-Gamma distribution in window *k* is modelled by the ploidy-dependent parameters 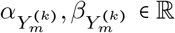, describing mean and dispersion, where 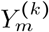 is the ploidy in the considered window. In each window, the estimated population frequencies serve as a proxy for the probability of sequenced reads. Note that the Poisson-Gamma distributions depend each on a ploidy level. This means that all windows assigned the same ploidy will refer to the same mean and dispersion parameters.

We propose a heuristic optimisation algorithm to automatically find the number of latent states of the HMM, and to assign them to the correct ploidy through the genotype likelihoods. Our implementation, described in the Supplementary Material, is a heuristic version of the well-known Expectation Conditional Maximisation (ECM) algorithm (Cappe et al., 2005).

### Simulated data

The required memory, runtime and ploidy detection power of `HMMploidy` were compared to the ones obtained by other methods using simulated data. We simulated sequencing reads under a wide range of scenarios using a previously proposed approach (Fumagalli, Vieira, Korneliussen, et al., 2013). Specifically, each locus is treated as an independent observation, without modelling the effect of linkage disequilibrium. The number of reads is modelled with a Poisson distribution with parameter given by the input depth multiplied by the ploidy level. At each locus, individual genotypes are randomly drawn according to a probability distribution defined by a set of population parameters (e.g., shape of the site frequency spectrum). Once genotypes are assigned, sequencing reads (i.e. nucleotidic bases) are sampled with replacement with a certain probability given by the base quality scores.

For comparing the performance of detecting ploidy between `HMMploidy` and existing tools, 100 simulations of M genomes are performed for every combination of ploidy (from 1 to 5, constant along each genome), sample size (1, 2, 5, 10, 20), and sequencing depth (0.5X, 1X, 2X, 5X, 10X, 20X). The sequencing depth is defined as the average number of sequenced bases at one site for each chromosomal copy (i.e. divided by the ploidy level). Each simulated genome has a length of 5Kb with all loci being polymorphic in the population.

Simulated data for the analysis of runtime and memory usage consist of 100 diploid genomes of length 10kb, 100kb, 1Mb, 10Mb. Each simulated genome comprises an expected proportion of polymorphic sites equal to 1%. The simulation scripts and pipelines are included in the Github and OSF repositories. Performance analysis was performed on a cluster node with four reserved cores of an Intel Xeon Gold 6130 @1.00GHz with 24GB of RAM and the Ubuntu 18.04.3 OS.

### Application to real data

To illustrate the use of `HMMploidy`, we apply it to sequencing data from 23 isolates of the pathogenic fungus *Cryptococcus neoformans* recovered from HIV-infected patients showing clinical evidence cryptococcal meningitis (Rhodes, Beale, Vanhove, et al., 2017). Whole-genome sequencing data was performed on an Illumina machine following an established protocol for sample preparation (Rhodes, Beale, and Fisher, 2014) and data processing (Rhodes, Beale, Vanhove, et al., 2017). Reads are mapped onto *C. neoformans* H99 reference genome (Loftus et al., 2005), yielding an average depth of approximately 100 reads per site. We generated an additional data set by randomly sampling only 20% of reads for each sample. All sequencing raw reads were retrieved from the European Nucleotide Archive under the project accession PRJEB11842.

## Results and discussion

### Predictive performance

We assess the power of `HMMploidy` to infer ploidy levels on simulated genomes ranging from haploid to pentaploid. Samples sizes varied from 1 to 20 individuals haplotypes, and sequencing depths from 0.5X to 20X. `HMMploidy` is compared to the two state-of-the-art methods `ploidyNGS` (Augusto Corrêa dos Santos et al., 2017) and nQuire (including a version with denoising option, `nQuire.Den`) (Weiß et al., 2018). The former performs a Kolmogorov-Smirnov test between the minor allele frequencies of the observed data and of simulated data sets at different ploidy levels (simulated at 50X). The latter models the minor allele frequencies with a Gaussian mixture model. We exclude depth-based methods because they are hardly applicable to low sequencing depth (Fig. S2, S3) and work as empirical visual checks rather than algorithmic procedures. While nQuire and `ploidyNGS` sweep the whole simulated genomes, `HMMploidy` analyses windows of 250bp, so the detection rate is calculated as the windows’ average, making the comparison deliberately more unfair to our method.

At low-depth (0.5X), `HMMploidy`’s power increases with sample size up to 20 - the largest we considered – in all scenarios excluding the tetraploid case (Fig. 2). This might be because it is difficult to distinguish diploid and tetraploid genotypes at such low depth. In the haploid and diploid case `ploidyNGS` has a remarkable 100% success at very low depths (Fig. 2). This is likely because having only few reads makes it easier to compare the data to a simulated genome with low ploidy level and a simpler distribution of observed alleles. However, this erratic behaviour disappears at higher ploidy levels, and `ploidyNGS` is generally outperformed by `nQuire.Den` and/or `HMMploidy`. `HMMploidy` is outperformed at low depth in the tetraploid scenario by both versions of nQuire. This might indicate that genotype likelihoods are not successful in modelling tetraploid genotypes as well as allele frequencies in this specific scenario.

**Figure 2. :**
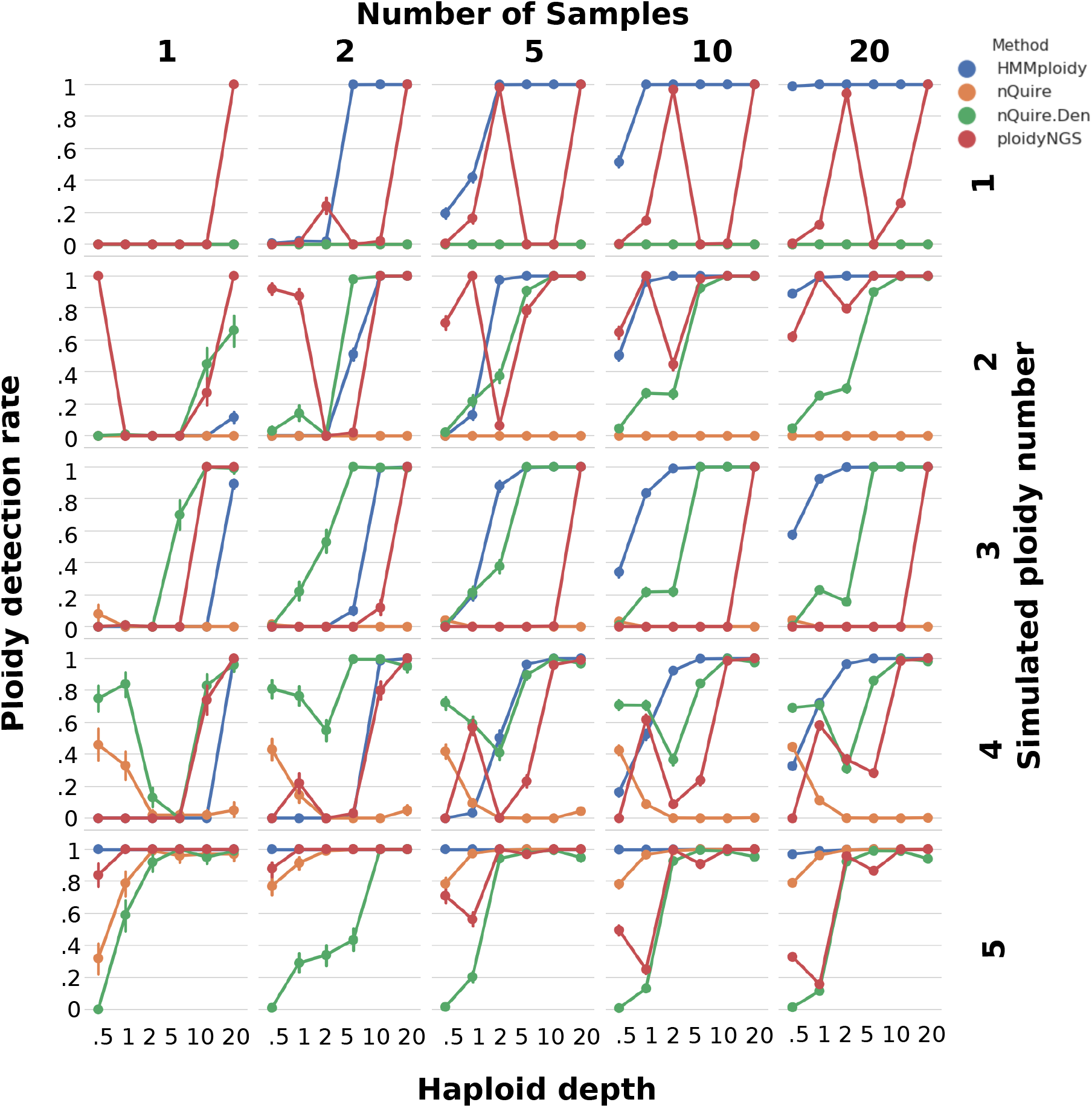
Comparison of ploidy detection rates for different methods at various experimental scenarios. The rate of detecting the correct ploidy (y-axis) is shown against the haploid sequencing depth (x-axis) for different sample sizes (on columns) and ploidy levels (on rows). For every simulated ploidy level, at each value of the sequencing depth we generate M genomes 100 times, where M is the number of simulated samples. The ploidy detection rate is the proportion of correctly detected ploidy levels in the genomic windows with the HMM method, and the proportion of correctly detected ploidy levels along each whole genome with the other tested methods.

Note also that none of the methods performs well with a single haploid sample. This happens because many loci show only one possible genotype, and even with the genotype likelihoods it is impossible to determine the multiplicity of the ploidy. With more samples it is possible to exploit loci with at least another allele to inform on the most likely genotype.

In all tested scenarios, `HMMploidy` greatly improves its accuracy with increasing sample size, with unique good performances at low depth (Fig. 2) not observed with other methods. Additionally, `HMMploidy` infers ploidy levels in sliding windows across the data (as in Fig. 3). Moreover, `HMMploidy` does not require a reference genome at a known ploidy, unlike `ploidyNGS`. `HMMploidy` can identify haploid genomes, unlike nQuire. Note that either deeper sequencing depth or larger sample size is likely to be required for `HMMploidy` to detect higher levels of ploidy, as the power of the method decreases with increasing ploidy (Fig. S4).

**Figure 3. :**
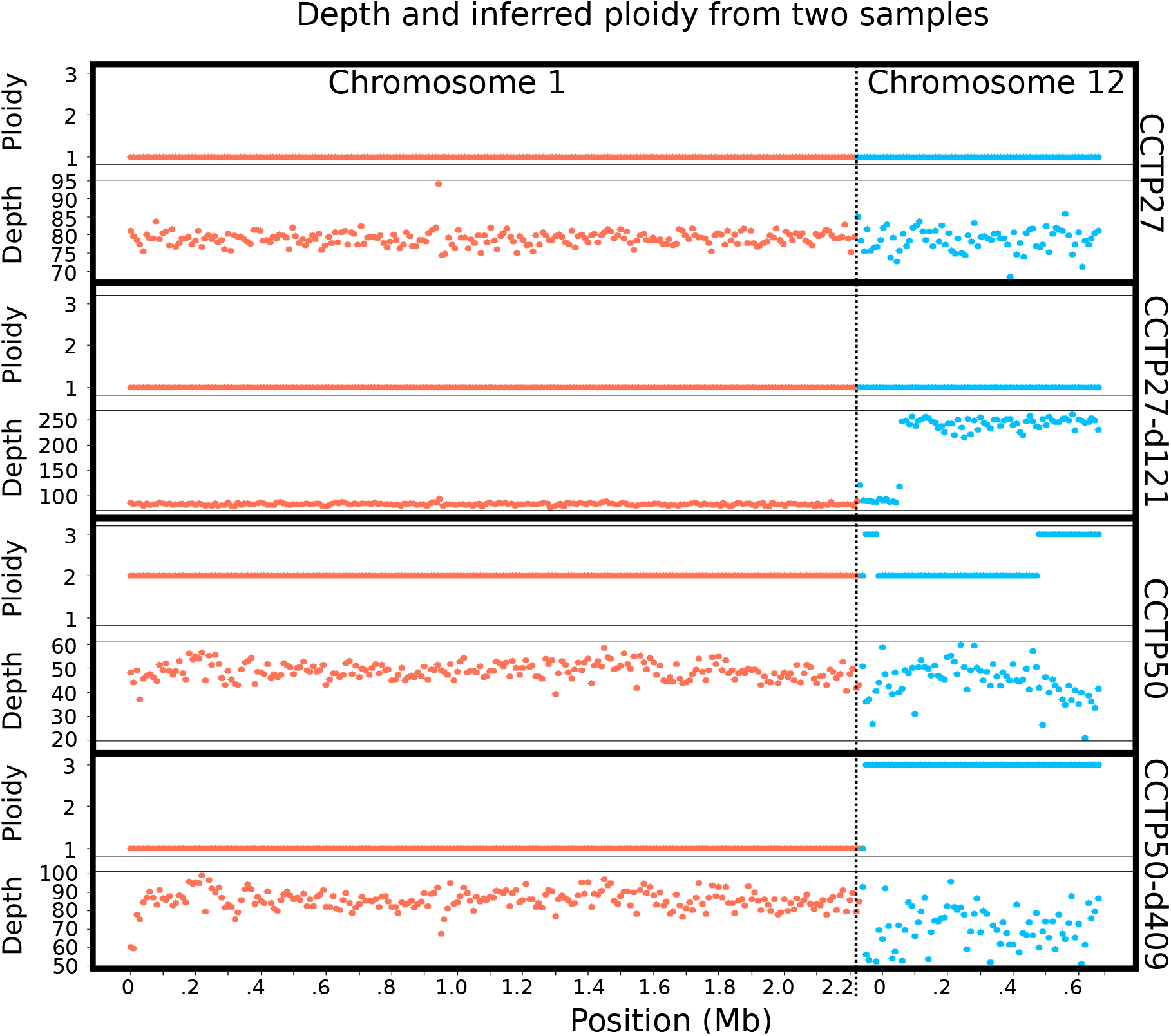
Inference of ploidy levels on two samples of *Cryptococcus neoformans* at different time points using. `HMMploidy`. Inferred ploidy and corresponding sequencing depth are shown in genomic windows for two samples at day 1 (CCTP27 and CCTP50), day 121 (CCTP27-d121) and 409 (CCTP50-d409) on chromosomes 1 and 12.

### Computational performance

The benchmark of `HMMploidy` shows a rather constant CPU time across genome lengths by keeping the number of windows fixed at *K* =100 (Fig. S5A). The shortest simulations are an exception, due to a very fast processing of the data to be used in the HMM. Occasionally, runtimes are elevated for cases where the inference algorithm is converging with difficulty. Fig. S5B shows the effect of increasing the number of windows on 10MB genomes. The growth of execution time follows linearly the increase of K, plus a probable extra overhead for preprocessing the data in many windows, showing that the forward-backward complexity *O* (∣𝒴∣^2^*K*) dominates the algorithm. In both the length-and windows-varying scenarios, memory usage was kept at an almost constant value of 350*MB*. This is possible thanks to the implementation of file reading and frequency estimation in C++. Both nQuire and `ploidyNGS` are obviously extremely fast and run in less than one second because they only need to calculate and compare observed allele frequencies, with a cost approximately comparable to the number of loci in the data. Therefore, their performance is not reported in the benchmark figures. Analogous trends on execution times would follow for genomes longer than 10MB and we expect `HMMploidy` to run without issues on larger genomes.

### Haploid depth

Note that `HMMploidy` trains a separate HMM on each genome even for larger sample sizes. As shown above, each HMM might require considerable CPU time if many windows are used, or if the heuristic ECM algorithm has a slow convergence. However, training a separate HMM on each genome allows the method to overcome two main issues: samples sequenced at different coverage, and ploidy varying among samples. When samples are sequenced at different coverage, it is common practice to standardise the sequencing depth across all genomes. However, this would make the estimation of the distributions of standardised counts difficult, especially in samples with noise, errors, and limited coverage. Additionally, two genomes could easily have two different ploidy levels matching the same distribution parameters. For example, a diploid-tetraploid sample where the two ploidy levels have observations’ mean parameters -1 and 1 could match haploid-diploid levels in another genome having the same mean parameters. The only case in which one can use the same HMM for all genomes is when they have all the same ploidy levels. However, this function is not implemented in `HMMploidy`. On the latter point, it would not be possible to detect sample-specific variation in ploidy levels when training the HMM on pooled genomic data. Therefore, training a separate HMM on each genome is an important feature in `HMMploidy`. However, a simple extension of `HMMploidy` would allow to estimate an HMM on the pooled data from multiple genomes, and to initiate HMM parameters and number of latent states to reduce the model estimation runtime. These options might be implemented in future versions of the software.

### Application to real data

We used `HMMploidy` to infer ploidy variation in 23 isolates of *Cryptococcus neoformans* recovered from HIV-infected patients (Rhodes, Beale, Vanhove, et al., 2017). By analysing variation in normalised sequencing coverage, Rhodes and coworkers identified extensive instances of aneuploidy, especially on chromosome 12, in several pairs of isolates (Rhodes, Beale, Vanhove, et al., 2017), in line with previous findings using karyotypic analysis (Ormerod et al., 2013). We sought to replicate these inferences using `HMMploidy` and assessed its performance on a downsampled data set to mirror data uncertainty.

In accordance with the original study (Rhodes, Beale, Vanhove, et al., 2017), we retrieve patterns of polyploidy and aneuploidy within each isolate. Most of the analysed samples are haploid (Fig. 3 and Fig. S6-S28). Interestingly, samples CCTP27 and CCTP27 at day 121 (CCTP27-d121) are inferred to have the same ploidy, even though CCTP27-d121 triplicates its sequencing depth on chromosome 12 (Fig. 3). We interpret this pattern as one CNV instance spanning most of chromosome 12 for CCTP27-d121. In fact, despite the increase in depth, the data is modelled as a haploid chromosome by the genotype likelihoods. This further illustrates the importance of jointly using information on genotypes and depth variation to characterise aneuploidy and CNV events. Sample CCTP50 had on average a higher depth at day 409, but chromosome 1 changed from diploid (day 1) to haploid (day 409). Chromosome 12 was triploid at day 409 although the high variability of sequencing depth is not informative on the ploidy.

Notably, we were able to retrieve the same patterns of predicted ploidy variation when artificially downsampling the sequencing data to 20% of the original data set (Fig. S6-S28). Interestingly, `ploidyNGS`, nQuire and `nQuire.Den` infer the highest tested ploidy in almost all windows of the 23 samples (Supplementary Table 1). This is likely because these methods fit the distribution of widely varying allele frequencies in each sample with the most complex ploidy model, as they do not consider the information of genotype likelihoods.

Cryptococcal meningitis, caused by the fungal yeasts *Cryptococcus neoformans* and *Cryptococcus gattii*, is a severe infection mostly affecting HIV/AIDS patients (May et al., 2016). Oral fluconazole antifungal therapies are widely used for treatment of Cryptococcal meningitis, although their efficacy is reported to be poor especially in Sub-Saharan Africa (Longley et al., 2008). Resistance to antifungal drugs is thought to be responsible for such poor outcomes and relapse episodes, but its molecular mechanisms are not yet understood (Stone et al., 2019). Resistance to oral fluconazole antifungal drugs in *Cryptococcus neoformans* was associated with aneuploidy (Sionov et al., 2013). Recent genomic studies identified multiple occurrences of aneuploidy in resistant and relapse isolates (Stone et al., 2019). Our genomics inferences of aneuploidy in *Cryptococcus neoformans* from HIV-infected patients can serve as diagnostic and molecular surveillance tools to predict and monitor drug resistance isolates, whilst further providing novel insights into the pathogen’s evolution (Rhodes, Desjardins, et al., 2017) We envisage that `HMMploidy` can be deployed to large-scale genomics data of pathogenic species to characterise aneuploidy-mediated drug resistance.

### Conclusions

Here we introduce `HMMploidy`, a method to infer ploidy levels suitable for low-and mid-depth sequencing data, as it jointly uses information from sequencing depth and genotype likelihoods. `HMMploidy` outperforms traditional methods based on observed allele frequencies, especially when combining multiple samples. We predict that `HMMploidy` will have a broad applicability in studies of genome evolution beyond the scenarios illustrated in this study. For instance, the statistical framework in `HMMploidy` can be adopted to infer aneuploidy in cancerous cells (Ben-David and Amon, 2020), or partial changes of copy numbers in polyploid genomes due to deletions or duplications (Vu et al., 2017).

## Acknowledgements

Preprint version 6 of this article has been peer-reviewed and recommended by Peer Community In Mathematical and Computational Biology (https://doi.org/10.24072/pci.mcb.100010). We are grateful to Alan Rogers, Barbara Holland, Benjamin Peter, Nicolas Galtier, and several anonymous reviewers for improving the manuscript. We wish to thank Anders Albrechtsen for providing feedback on the formulation of genotype likelihoods. We would like to thank GenomeDK and Aarhus University for providing computational resources and support.

## Fundings

SS was supported by a grant from Novo Nordisk Foundation (NNF20OC0063268). JR and MCF were supported by a grants from Natural Environmental Research Council (NERC; NE/P001165/1 and NE/P000916/1), the UK Medical Research Council (MRC; MR/R015600/1) and the Wellcome Trust (219551/Z/19/Z). MCF is a CIFAR Fellow in the ‘Fungal Kingdom’ programme. We acknowledge support from the Erasmus+ programme to IA.

## Conflict of interest disclosure

The authors declare that they comply with the PCI rule of having no financial conflicts of interest in relation to the content of the article. The authors declare the following non-financial conflict of interest: Matteo Fumagalli and François Balloux are recommenders of PCI.

## Data, script and code availability

Data are available online: https://doi.org/10.17605/OSF.IO/5F7AR

Script and codes are available online: https://doi.org/10.5281/zenodo.7116023

## Supplementary information availability

Supplementary information is available online: https://doi.org/10.1101/2021.06.29.450340

## Notes

https://doi.org/10.17605/OSF.IO/5F7AR

https://doi.org/10.5281/zenodo.7116023

### Competing Interest Statement

The authors have declared no competing interest.

### Summary of Updates

DOI links have been provided to external data and supplementary information.

