## Supplementary Information for "HMMploidy: inference of ploidy levels from short-read sequencing data"

### 1 Supplementary Material

#### 1.1 Supplementary Methods

This section details the methods used for the implementation of `HMMploidy`. We assume that $O = (O_1, \ldots, O_M)$ is the observed NGS data for $M$ sequenced genomes at $N$ loci. For each $m$-th genome and $n$-th locus we define $Y_{mn}$, $G_{mn}$ and $O_{mn}$ as the ploidy, genotype and sequencing data, respectively. $O_{mn}$ consists of $C_{mn}$ nucleotides from aligned reads. $G_{mn}$ takes values in $\{0, 1, ..., Y_{mn}\}$, where the numbers denote the amount of derived alleles assuming that the site is a diallelic SNP.

#### 1.2 Genotype likelihood for arbitrary ploidy number

We review here the concept of genotype likelihood as previously and extensively described (33; 1; 40). Genotype likelihoods are calculated using the base quality of each nucleotide, and here we extend the concept to an arbitrary ploidy. Genotype likelihoods are at the core of `HMMploidy`, because they are used to assign a probability of observing nucleotides at each locus given a possible genotype. Calculating genotype likelihoods for each ploidy (which in turn has its own set of genotypes) allows `HMMploidy` to obtain a set of likelihoods for each nucleotide locus given a ploidy's possible genotypes.

Given $m, n$, consider the $C_{mn}$ aligned nucleotides observed in $O_{mn}$, and the ploidy $Y_{mn}$. Let $O_{mnc}$ be the $c$-th nucleotide at locus $n$, and $G_{mn} = a_1 + \cdots + a_{Y_{mn}}$. Here, both $O_{mnc}$ and the $a_i$'s take value in $\{0, 1\}$, so that the sum of the $a_i$'s defines the genotype. Values $\{0, 1\}$ can be considered as the reference and alternate allele, respectively, with other polarisations being possible (e.g., ancestral and derived allele). Nucleotides and their errors across reads at locus $n$ are assumed to be independent (33) allowing the genotype likelihood to be written as a product across nucleotides in the following form:

$$L(G_{mn}|O_{mn}, Y_{mn}) = p(O_{mn}|G_{mn}) = \prod_{c=1}^{C_{mn}} p(O_{mnc}|G_{mn}).$$

Assuming parental alleles are randomly sampled, the probability of observing the $c$-th nucleotide given the observed genotype is calculated as

$$p(O_{mnc}|G_{mn}) = \frac{1}{Y_{mn}} \sum_{y=1}^{Y_{mn}} p(O_{mnc}|a_y).$$

Let $\epsilon_{mnc}$ be the probability of sequencing error calculated from the Phred quality score (15) for each observed nucleotide $O_{mnc}$. Then the $y$-th element of the above sum is modelled as

$$p(O_{mnc}|a_y) = \begin{cases} 1 - \epsilon_{mnc}, & \textit{if } O_{mnc} = a_y \\ \frac{\epsilon_{mnc}}{3} & \textit{otherwise} \end{cases}$$

This calculation of genotype likelihoods for polyploid genomes can be considered as an improved generalisation of (8) without relying on their assumption of constant error rate occurring only between the reference and alternate alleles.

##### 1.3 Estimation of population frequencies

Population allele frequencies are calculated prior to the HMM optimisation to decrease the computational time. Specifically, the population frequency $F_n$ at the $n$-th locus is estimated under the assumption of ploidy level being arbitrarily very high to let frequencies represent any possible genotype.

Let $\hat{F}_{mn}$ be the observed minor allele frequency for sample $m$ at locus $n$. The population frequency estimator for $F_n$, say $\hat{F}_n$, is defined as

$$\hat{F}_n = \frac{1}{C_n} \sum_{m=1}^{M} C_{mn} \hat{F}_{mn}, \tag{1}$$

where $C_n = \sum_{m=1}^{M} C_{mn}$.

##### 1.4 Hidden Markov Model for ploidy inference

Here, the HMM is defined, and the inferential process of ploidy levels from the HMM is illustrated. Mathematical details, proofs and algorithm analysis of the HMM for ploidy inference (Fig. S1) are presented here.

The Markov chain of ploidy levels is characterised by a $|\mathcal{Y}| \times |\mathcal{Y}|$ transition matrix $\boldsymbol{A}$, and a $|\mathcal{Y}|$-long vector $\boldsymbol{\delta}$ of starting probabilities for the first latent state. Here, $\mathcal{Y}$ is the set of ploidy levels included in the model, and $|\mathcal{Y}|$ is its cardinality. The average depth for genome $m$ in window $k$ is characterised by the ploidy-dependent parameters $\alpha_{Y_m^{(k)}}, \beta_{Y_m^{(k)}} \in \mathbb{R}$, describing mean and dispersion of the data, for each $Y_m^{(k)} \in \mathcal{Y}$. For brevity we write the parameters of the depth distribution in vector form, i.e. $\boldsymbol{\alpha}$, $\boldsymbol{\beta}$. The allele frequencies calculated through Equation (1) in the $k$-th window of loci serve as a proxy for the probability of sequenced reads.

**Heuristic Expectation Conditional Maximization (HECM)** The ECM algorithm is used to infer the parameters $\boldsymbol{A}$, $\boldsymbol{\delta}$ modelling the sequence of ploidy levels. This is done in two iterative steps by exploiting the ploidy-dependent distributions of the observed data (sequenced reads and coverage in each window). The first step is the well-known forward-backward algorithm (39; 9), that computes in each window the probability of a ploidy given all the observed data. This is done in an efficient way through dynamic programming and exploitation of Markov properties with computational complexity $\mathcal{O}(|\mathcal{Y}|^2 K)$ (i.e. linear in the number of loci windows) by implementing two calculation sweeps, starting respectively at the end and at the beginning of the observation sequence.

The forward-backward algorithm thus creates the mathematical link between ploidy levels and observed data, and allows us to update the parameters governing the Markov chain of ploidy levels with the ECM algorithm in a subsequent step. The ECM algorithm maximizes a value (called intermediate quantity) strictly related to the likelihood of the model, where the free variables of the maximization are the matrix $\boldsymbol{A}$, the vector $\boldsymbol{\delta}$ and the parameters of the distribution of observed data. This procedure continues iteratively by recalculating the forward-backward posteriors and the update parameters with the ECM, until the intermediate quantity cannot be further improved.

We start by illustrating the steps of the ECM algorithm, and subsequently adding the heuristic procedure. For ease of notation, $\lambda$ is the tuple of parameters $(\boldsymbol{A},\boldsymbol{\delta},\boldsymbol{\beta},\boldsymbol{\alpha}) \in \Lambda$, considered as two separate tuples: $\lambda = (\lambda_1,\lambda_2) = ((\boldsymbol{A},\boldsymbol{\delta},\boldsymbol{\beta}),(\boldsymbol{\alpha})) \in \Lambda_1 \times \Lambda_2$. Given the parameters $\lambda^{\ell-1}$ calculated at the $(\ell-1)$-th step of the ECM, the $\ell$-th iteration to calculate $\lambda^{\ell}$ follows the steps below:

1. calculate the intermediate quantity

$$Q(\lambda_1^{\ell}|\lambda^{\ell-1}) = \mathbb{E}\Big[\ln p\Big(O_m^{(1:K)}, Y_m^{(1:K)}|(\lambda_1^{\ell},\lambda_2^{\ell-1})\Big)\,\Big|\,O_m^{(1:K)},\lambda^{\ell-1}\Big];$$

2. calculate $\lambda_1^{\ell} = \arg_{\lambda_1^{\ell}\in\Lambda_1}\max Q(\lambda_1^{\ell}|\lambda^{\ell-1}))$;
3. calculate the intermediate quantity $Q(\lambda_2^{\ell}|(\lambda_1^{\ell},\lambda_2^{\ell-1}))$ analogously to step 1;
4. calculate $\lambda_2^{\ell} = \arg_{\lambda_2^{\ell}\in\Lambda_2}\max Q((\lambda_2^{\ell})|(\lambda_1^{\ell},\lambda_2^{\ell}))$.

Here, we used $O_m^{(1:K)}$, $Y_m^{(1:K)}$ to denote $O_m^1,\dots,O_m^K$ and $Y_m^1,\dots,Y_m^K$, respectively.

The ECM algorithm for a HMM with negative binomial observations thus consists of two EC-steps and two maximization steps. Specifically for the four steps above:

1. the first EC-step calculates the expected complete-data log-likelihood with the Markov chain parameters and the dispersion ($\boldsymbol{\beta}$) parameters unknown and to be estimated at the next M-step (maximization step), conditionally to the mean ($\boldsymbol{\alpha}$) parameters estimated at the previous iteration of ECM;
2. the first M-step maximizes the intermediate quantity calculated at the first step w.r.t. the unknown parameters;
3. the second EC-step replicates the first one inverting the roles of known and unknown parameters;
4. the second intermediate quantity can be maximized w.r.t. the mean parameters.

The EC-step of the ECM algorithm is very similar to the classical forward-backward formulation in the E-step of the EM algorithm (7; 39). The E-step expresses the expected complete-data log-likelihood with the all HMM parameters unknown and to be estimated at the next M-step. The E-step works for observations distributed with one parameter, or multiple parameters whose maximization equations can be solved in normal form, i.e. by isolating the parameter of interest in each equation (e.g. Poisson and Gaussian distribution). The EC-step is a formulation of the E-step where only a portion of the HMM parameters can be estimated in one maximization step (the M-step). This is a characteristic of emission distributions whose parameters can be estimated only in function of each other in a system of equation (e.g. gamma and negative binomial distributions). The average depths are modelled with a negative binomial distribution to take data overdispersion into account (12).

The calculation of $\boldsymbol{A}, \boldsymbol{\delta}$ at iteration $\ell$ is solved by using the classical forward-backward algorithm (39; 9), therefore we will only briefly mention the necessary elements of it, while we analyzed more in depth the estimation of means and dispersions.

The scope of each ECM iteration is to maximize the intermediate quantities to achieve the highest value of the complete data log-likelihood. In this way, at each iteration, new parameters can be used to rewrite $Q$ and remaximize it until convergence. It is worth remembering again that the resulting parameters maximize a quantity different from the log-likelihood of the observed data - the ECM uses two forms of $Q$ to make the maximization feasible, since expressing the log-likelihood directly is not concretely achievable.

The intermediate quantity can be explicitly written as the sum of three terms involving separately the matrix $\boldsymbol{A}$, the vector $\boldsymbol{\delta}$ and the vectors $\boldsymbol{\alpha}, \boldsymbol{\beta}$:

$$Q(\lambda_1^\ell|\lambda^{\ell-1}) = \sum_{Y_m^{(1:K)} \in \mathcal{Y}} \ln(\delta^\ell_{Y_m^{(1)}}) p(Y_m^{(1:K)}|O_m^{(1:K)}, C_m^{(1:k)}, \lambda^{\ell-1}) \tag{2}$$

$$+ \sum_{Y_m^{(1:K)} \in \mathcal{Y}} \textstyle\sum_{k=2}^K \ln(\boldsymbol{A}^\ell_{Y_m^{(k-1)} Y_m^{(k)}}) p(Y_m^{(1:K)}|O_m^{(1:K)}, C_m^{(1:k)}, \lambda^{\ell-1}) \tag{3}$$

$$+ \sum_{Y_m^{(1:K)} \in \mathcal{Y}} \sum_{k=1}^K \left( \ln\left(p(O_m^{(k)}|Y_m^{(k)}, F^{(k)})\right) + \ln\left(p(C_m^{(k)}|Y_m^{(k)}, \alpha^{\ell-1}_{Y_m^{(k)}}, \beta^\ell_{Y_m^{(k)}})\right)\right)$$

$$p(Y_m^{(1:K)}|O_m^{(1:K)}, C_m^{(1:k)}, \lambda^{\ell-1}) \tag{4}$$

where the logarithm of a matrix is intended element-wise.

Consider the $(m,k)$-th forward variable defined by

$$f(y_m^{(k)}) = p(O_m^{(1:k)}, C_m^{(1:k)}, Y_m^{(k)} = y_m^{(k)}|\lambda),$$

that is, the probability of the first $k$ observations and $k$-th ploidy $y_m^{(k)}$ given the parameters $\lambda$. Define the $(m,k)$-th backward variable as

$$b(y_m^{(k)}) = p(O_m^{(k+1:K)}, C_m^{(k+1:K)}|Y_m^{(k)} = y_m^{(k)}, \lambda),$$

that is, the probability of the latest $(K-k)$ observations, given the $k$-th ploidy $y_m^{(k)}$ and the parameters $\lambda$. The forward and backward variables can be computed with an iterative procedure (39, eq. 19,20,24,25) and allow us to calculate efficiently the likelihood of the data as

$$p(O_m^{(1:K)}, C_m^{(1:k)}|\lambda) = \sum_{y_m^{(k)} \in \mathcal{Y}} f(y_m^{(k)}) b(y_m^{(k)}) \quad \text{for any } k = 1, \ldots, K.$$

The two terms in the equation lines (2) and (3) include only the parameters $\boldsymbol{\delta}$ and $\boldsymbol{A}$, respectively. This simplifies finding optimisation formulae for those parameters by considering separately each term of lines (2) and (3). Such optimisation equations for $\boldsymbol{\delta}$ and $\boldsymbol{A}$ are easily derived through Lagrange multipliers (39, eq. 40a,40b). This does not solve the second step of the ECM algorithm, because the optimum for $\boldsymbol{\beta}$ is still not calculated.

It is easy to see that both $\boldsymbol{\alpha}$ and $\boldsymbol{\beta}$ concur in defining line (4). This is what originates the conditional nature of the ECM algorithm, i.e. $\boldsymbol{\alpha}$ and $\boldsymbol{\beta}$ cannot be optimised independently. Therefore we first optimise $\boldsymbol{\beta}$ considering the values of $\boldsymbol{\alpha}$ calculated at the $(\ell-1)$-th iteration of the ECM algorithm. Using the forward and backward variables, and excluding terms independent from the Poisson-Gamma parameters, Equation (4) can be written as follows:

$$\begin{aligned}
&\sum_{Y_m^{(1:K)}\in\mathcal{Y}} \sum_{k=1}^{K} u(m,k) \ln\left(p(C_m^{(k)}|Y_m^{(k)}, \alpha^{\ell-1}_{Y_m^{(k)}}, \beta^{\ell}_{Y_m^{(k)}})\right) \\
&= \sum_{Y_m^{(1:K)}\in\mathcal{Y}} \sum_{k=1}^{K} u(m,k) \ln\left(\frac{\Gamma(\alpha^{\ell-1}_{Y_m^{(k)}} + C_m^{(k)})}{\Gamma(C_m^{(k)}+1)\Gamma(\alpha^{\ell-1}_{Y_m^{(k)}})}\right) \\
&+ \sum_{Y_m^{(1:K)}\in\mathcal{Y}} \sum_{k=1}^{K} u(m,k)\left(C_m^{(k)} \ln\left(\frac{1}{\beta^{\ell}_{Y_m^{(k)}}+1}\right) + \alpha^{\ell-1}_{Y_m^{(k)}} \ln\left(\frac{\beta^{\ell}_{Y_m^{(k)}}}{\beta^{\ell}_{Y_m^{(k)}}+1}\right)\right)
\end{aligned}$$

where $u^{\ell}(m,k) = f(y_m^{(k)})b(y_m^{(k)})/p(O_m^{(1:K)}, C_m^{(1:k)}|\lambda^{\ell})$. By setting the partial derivative of $Q(\lambda_1^{\ell}|\lambda^{\ell-1})$ w.r.t. a certain $\beta^{\ell}_{y_m^{(k)}}$, $y_m^{(k)} \in \mathcal{Y}$ equal to zero as below, we will be able to calculate the optimum for the derivative's parameter:

$$\frac{\partial Q(\lambda_1^{\ell}|\lambda^{\ell-1})}{\partial \beta^{\ell}_{y_m^{(k)}}} = \sum_{k=1}^{K} -u(m,k)\frac{C_m^{(k)}}{\beta^{\ell}_{y_m^{(k)}}+1} + \sum_{k=1}^{K} u(m,k)\frac{\alpha^{\ell-1}_{Y_m^{(k)}}}{\beta^{\ell}_{y_m^{(k)}}\left(\beta^{\ell}_{y_m^{(k)}}+1\right)} = 0.$$

Solving for $\beta^{\ell}_{y_m^{(k)}}$ leads to the optimum of the parameter:

$$\beta^{\ell}_{y_m^{(k)}} = \frac{\alpha^{\ell-1}_{Y_m^{(k)}} \sum_{k=1}^{K} u(m,k)}{\sum_{k=1}^{K} u(m,k) C_m^{(k)}}.$$

This completes the step 2 of the ECM. In our implementation of `HMMploidy`, we want to leverage the information contained in the genotype likelihoods, whose partial derivative goes to zero and in principle are not integrated in the optimisation. In `HMMploidy`, we add the genotype likelihoods to the depth distribution prior to optimisation, so that forward and backward variables contain information on both depth and genotypes, and allow the identification of different states with distinct ploidy levels.

The value of $Q(\lambda_2^{\ell}|(\lambda_1^{\ell}, \lambda_2^{\ell-1}))$ can be easily calculated as in step 1, and by setting the partial derivative of $Q(\lambda_2^{\ell}|(\lambda_1^{\ell}, \lambda_2^{\ell-1}))$ w.r.t. $\alpha^{\ell}_{y_m^{(k)}}$, $y_m^{(k)} \in \mathcal{Y}$, to zero, we obtain:

$$\sum_{k=1}^{K} u(m,k)\left( \ln\Big(\frac{\beta^{\ell}_{y_m^{(k)}}}{\beta^{\ell}_{y_m^{(k)}}+1}\Big) + \psi_0\big(\alpha^{\ell}_{y_m^{(k)}} + C_m^{(k)}\big) - \psi_0\big(\alpha^{\ell}_{y_m^{(k)}}\big)\right) = 0.$$

Solving for $\alpha^{\ell}_{y_m^{(k)}}$ is done through the Newton-Rapson method (7), completing step 4 of the ECM.

**Heuristic step and ploidy inference** The ECM algorithm is repeated as an iterative sequence of EC and M steps, until the expected conditional log-likelihood of the model satisfies a convergence criteria. When convergence is achieved, `HMMploidy` performs the heuristic step, by running few iterations of the ECM over the HMM, where the set of ploidy levels is reduced by one, and the parameters for initialisation are the final ones from the ECM. We assume that, if the HMM has an overfitting set of ploidy levels, observation parameters are overlapping (27) for two or more ploidy levels. Therefore, removing one unnecessary ploidy requires only few extra iterations for the EM to converge again. The Bayesian Information Criterion (BIC) (7; 9) is used to compare the HMM with the reduced HMMs. If there is a reduced HMM with a better BIC score, then the ECM runs again on such HMM, otherwise it stops. Such method is an adaptation of the suggestion in (27).

After the HMM is reduced through the BIC comparison, we reduce the transition matrix between ploidy levels, i.e. we remove ploidy values for which there is almost zero probability of lasting a reasonable number of adjacent windows. In other words, we remove ploidy levels that will last for the length of one or few more windows of loci. Once the HMM parameters are determined through the heuristic sweep, the standard Viterbi algorithm (48) is applied to infer the most likely sequence of ploidy levels from the parameters of the HMM. The Viterbi algorithm is another example of dynamic programming, allowing to bypass the calculation of all possible $|\mathcal{Y}|^K$ sequences of ploidy levels to determine which one is the best.

**Reduction of the transition matrix** An important element of a HMM is the transition matrix between states and the meaning of each state. Thanks to the heuristic ECM, `HMMploidy` is able to assign a ploidy to each state of the Markov chain in an unsupervised mode without overfitting the data. However, one needs to check whether transitions between states follow a biological meaning. For example, it is unlikely that a ploidy occurs only in a small window of loci, and then shifts again to the previous value, i.e. such event is likely due to noise or other biological artefact altering the quality and behaviour of the data (e.g. the presence of a centromere).

Once the HECM algorithm has converged to a set of parameters $\lambda \in \Lambda$, it is possible to perform an optional filtering on the transition matrix $\boldsymbol{A}$ of the HMM. Given the matrix $\boldsymbol{A}$ of size $|\mathcal{Y}| \times |\mathcal{Y}|$, the time of permanence in a state

$y \in \mathcal{Y}$ has geometric distribution with parameter $\boldsymbol{A}_{yy}$ (10). If the user expects that a ploidy level has to remain uninterrupted for at least a certain number of windows $N$, then a corresponding minimum value for the parameter of the geometric distribution can be estimated. In fact, the probability of permanence in ploidy $y$ for at least $N > 0$ windows is given by the cumulative distribution function of the geometric distributions, that is, $1 - (1 - y)^N$.

Given $N$, `HMMploidy` calculates the minimum value of $y$ that has to be on the diagonal of $\boldsymbol{A}$. Rows and columns corresponding to diagonal entries lower than $y$ are cancelled and $\boldsymbol{A}$ is rendered stochastic again. Corresponding values of $\delta$, $\alpha$, $\beta$ are also removed. Afterwards, the HMM optimisation is performed again on the new subset of ploidy levels for adjustment of the remaining parameters.

**Application in presence of sparse polymorphic sites** Given an individual $m$, consider each $k$-th window of its genome. In presence of very few polymorphic sites in each window, the genotype likelihoods might not be enough to determine the ploidy, especially when the data is at low-depth and in presence of error, as it is often the case with high-throughput data.

To consider this case, the option `useGeno` is added to the software. When `useGeno='yes'`, the HMM infers the ploidy numbers as explained in the main text. Otherwise, if When `useGeno='no'`, only the sequencing depth data is used to infer the hidden states of a binomial HMM initially. This allows to consider the largest possible windows of loci. Each latent state is then assigned a ploidy by maximising Equation (1) over all the windows with same hidden state.

##### 1.5 Results from the analysis of *Cryptococcus Neoformans*

Here we present all the inferred ploidy levels from the 23 isolates of *Cryptococcus Neoformans* from the original study (42). Each figure contains:

- In the first line, inferred ploidy levels from chromosome 1 and 12 using the full data,
- In the second line, inferred ploidy levels from chromosome 1 and 12 using 20% of the original sequencing data.

Most of the results from the downsampled data coincide with the inference from the whole data. Higher ploidy levels can be hard to detect in some cases, and are occasionally detected as a constant lower ploidy or as a highly varying sequence of adjacent ploidy levels. However, downsampling seems to recover a constant haploid chromosome 12 in sample cctp50 (Fig. S12B-D) according to what the sequencing depth indicates. This means that downsampling might reduce the effect of noisy data points that could alter the detected ploidy. In fact, the triploid sections of chromosome 12 are at the extremities of the chromosome, where the data is more affected by noise and in general by a lower sequencing quality.

All the other samples recover successfully the original ploidy levels in downsampled data. However, note that there are few changes in ploidy probably due to noise or the presence of reads close to the centromere (Fig. S21, S16, S15, S13, S14). Ploidy levels for *Cryptococcus Neoformans* samples inferred by competing methods `nQuire`, `nQuire.Den`, and `ploidyNGS` are presented in Supplementary Table 1.

#### 1.6 Supplementary Figures

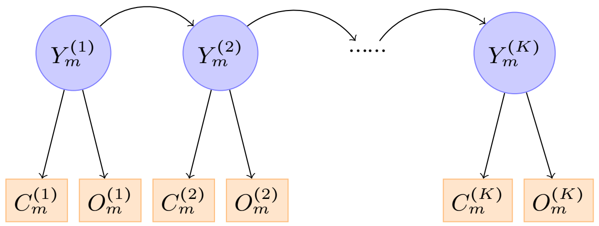


Figure S 1**: Hidden Markov Model for ploidy inference.** Graphical representation of the HMM to infer the ploidy levels of the $m$-th genome. $Y_m^{(k)}$ is the ploidy of the $k$-th window of genome $m$. The ploidy-dependent emissions consist of the average sequencing depth $C_m^{(k)}$ and the sequenced data $O_m^{(k)}$, whose distributions are respectively described by a Poisson-gamma distribution and by Equation (1).

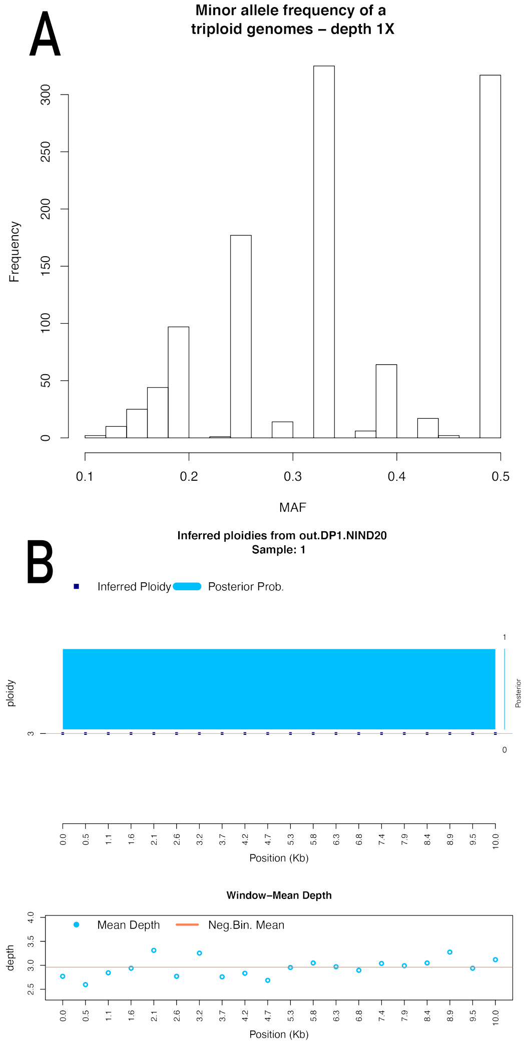


Figure S 2**: Histograms of minor allele frequencies and inferred ploidy with `HMMploidy` at low depth.** (A) Distribution of the minor allele frequencies of one simulated triploid genomes (out of a sample of 20 individuals) of 10kbp at depth $1X$ for the haploid state. It is not trivial to determine the ploidy by visual inspection of this graph. (B) Inferred ploidy with `HMMploidy` from the same individual on windows of 500 bases. Using the information contained in all the other individuals, it is possible to infer the correct ploidy.

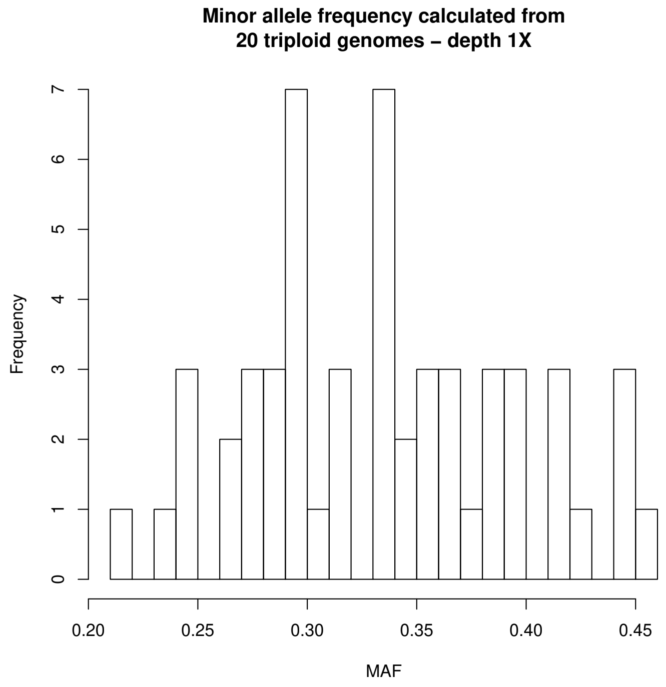


Figure S 3: **Histograms of minor allele frequencies for many samples at low depth.** Histogram of the estimated minor allele frequency for 20 simulated triploid individuals in a window of 500 bases. The distribution is closer to the one expected for a triploid individual, but it is still not possible to infer the ploidy by a simple visual inspection of the graph. The use of genotype likelihoods in `HMMploidy` supplies additional information to infer the correct ploidy.

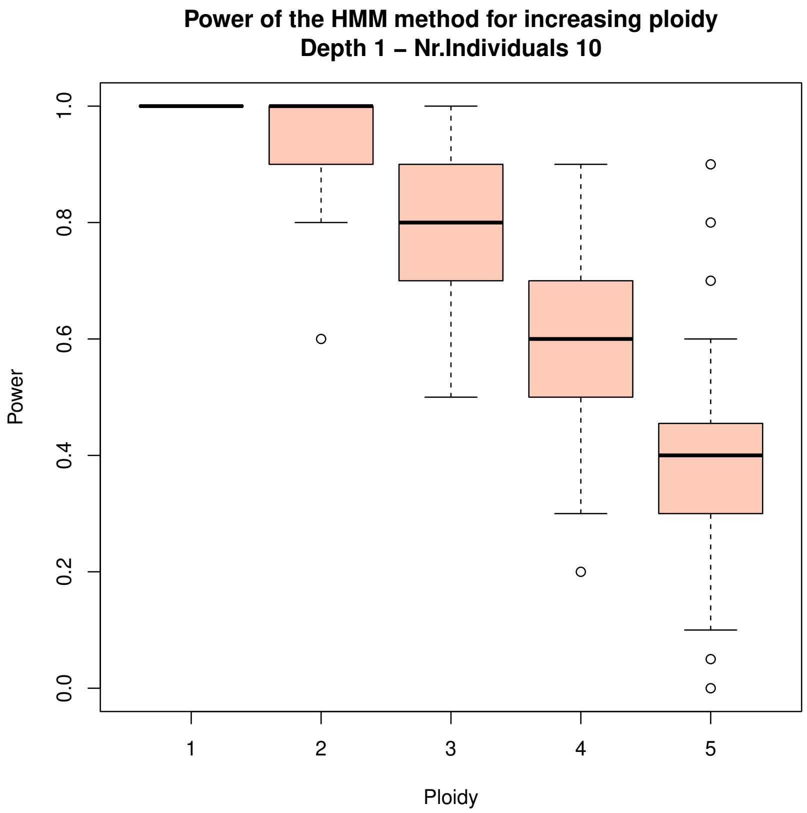


Figure S 4: **Relationship between ploidy levels and detection rate.** Power of `HMMploidy` to detect the correct ploidy level (on y-axis) on simulated genomes with increasing ploidy (on x-axis) from one to six at depth $1X$. The power decreases with higher ploidy numbers because genotype likelihoods lack information to characterise correct genotypes.

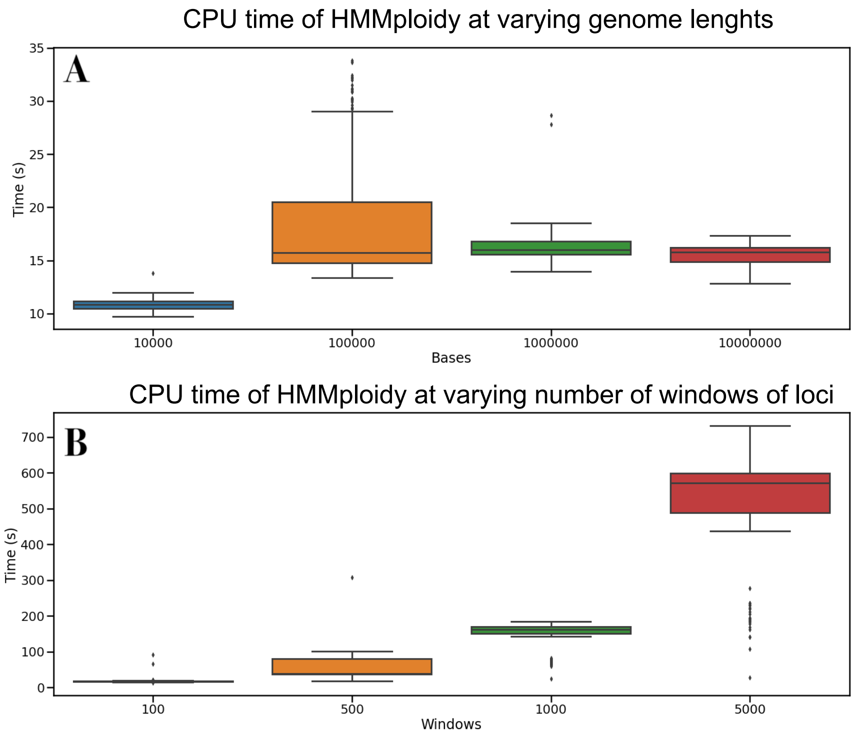


Figure S 5: **CPU running time for `HMMploidy`.** (A) CPU running time of `HMMploidy` by simulating genomes of various lengths and keeping the windows number to 100. The time is quite constant, meaning that the loading and processing of the data is very fast, and most of the time is taken by the HMM inference. (B) CPU running time of `HMMploidy` by increasing the number of windows on a $10MB$ genome. The time grows accordingly with K in an almost linear fashion (due to a probable overhead for preprocessing the data in many windows), as predicted by the computational cost of the forward-backward algorithm.

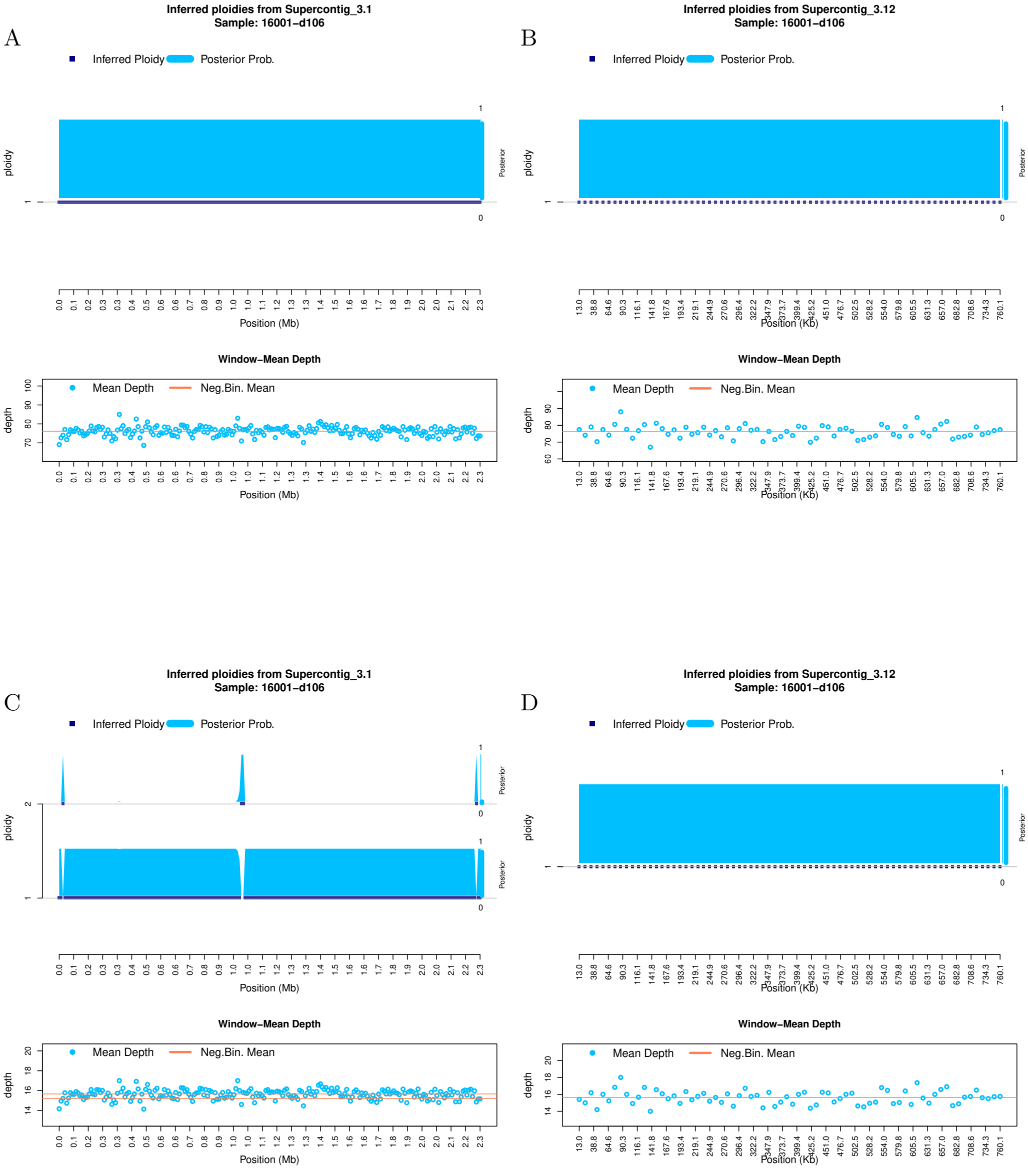


Figure S 6**: Ploidy inference on full and downsampled sequencing data.** Inferred ploidy levels from `HMMploidy` for chromosome 1 and 12 of isolate 16001-d106. **(A-B)** Results using the whole data on chromosomes 1 (A) and 12 (B). **(C-D)** Results using the data downsampled to 20% of its original depth on chromosomes 1 (C) and 12 (D).

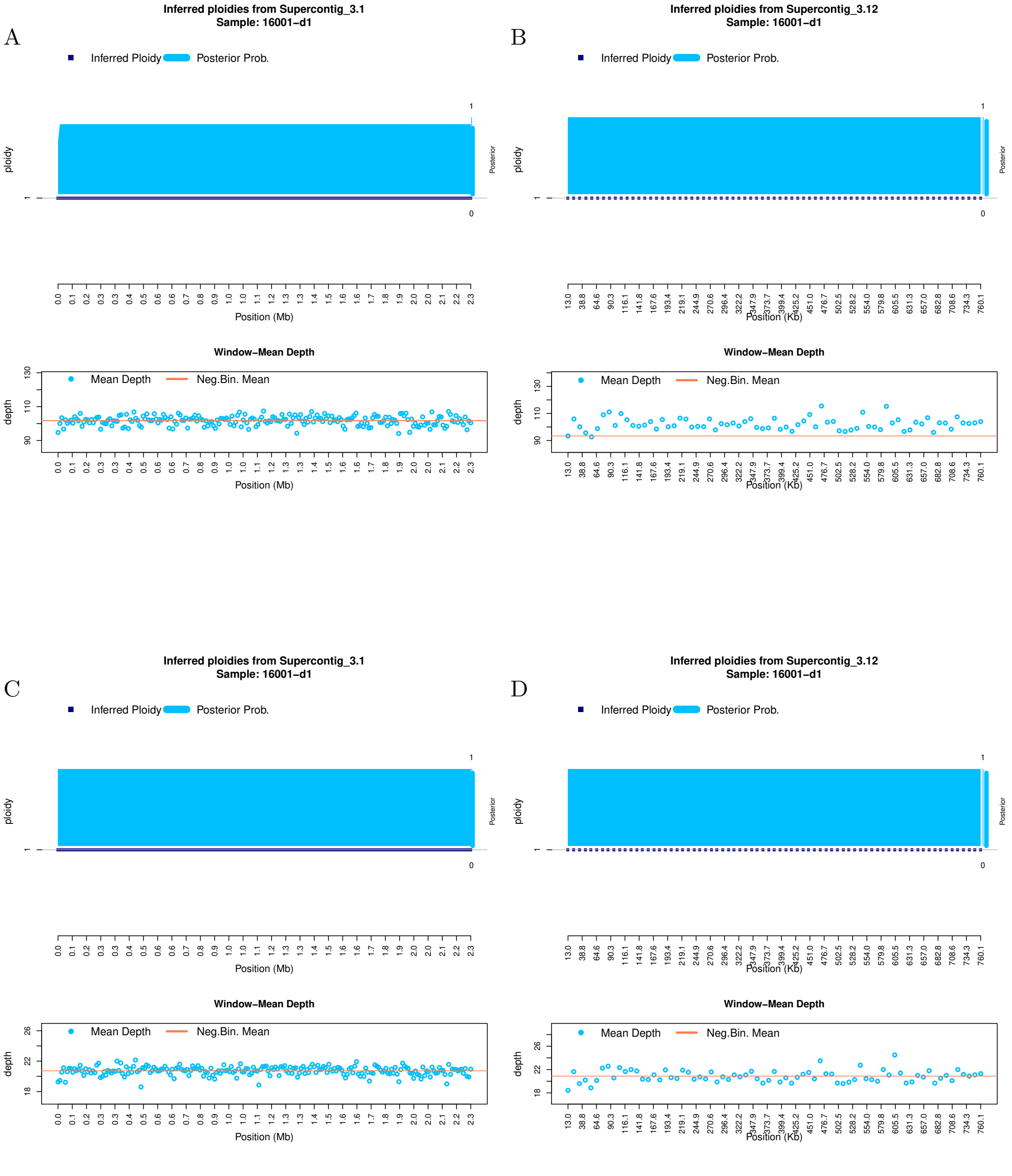


Figure S 7**: Ploidy inference on full and downsampled sequencing data.** Inferred ploidy levels from `HMMploidy` for chromosome 1 and 12 of isolate 16001-d1. **(A-B)** Results using the whole data on chromosomes 1 (A) and 12 (B). **(C-D)** Results using the data downsampled to 20% of its original depth on chromosomes 1 (C) and 12 (D).

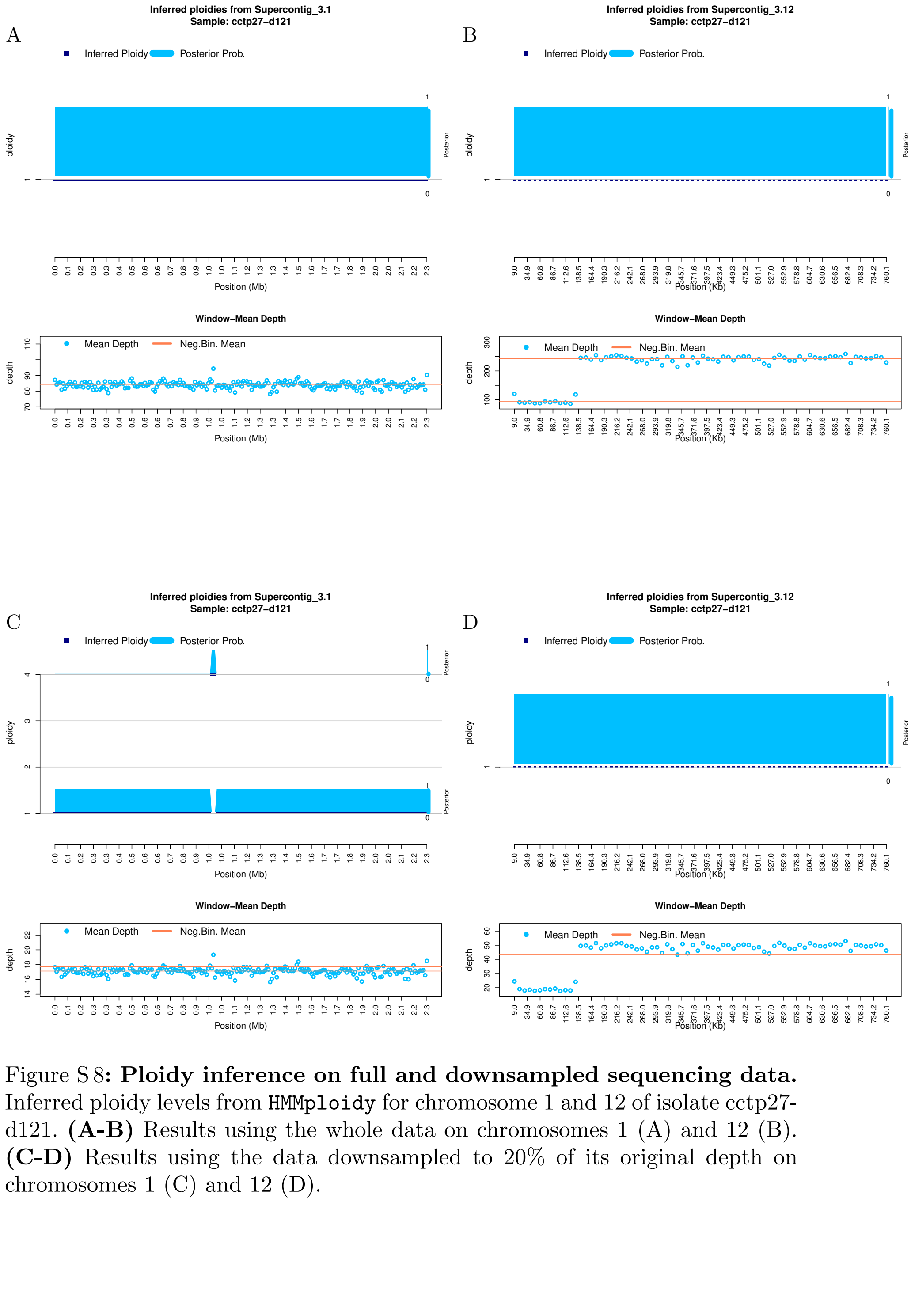


Figure S 8**: Ploidy inference on full and downsampled sequencing data.** Inferred ploidy levels from `HMMploidy` for chromosome 1 and 12 of isolate cctp27-d121. **(A-B)** Results using the whole data on chromosomes 1 (A) and 12 (B). **(C-D)** Results using the data downsampled to 20% of its original depth on chromosomes 1 (C) and 12 (D).

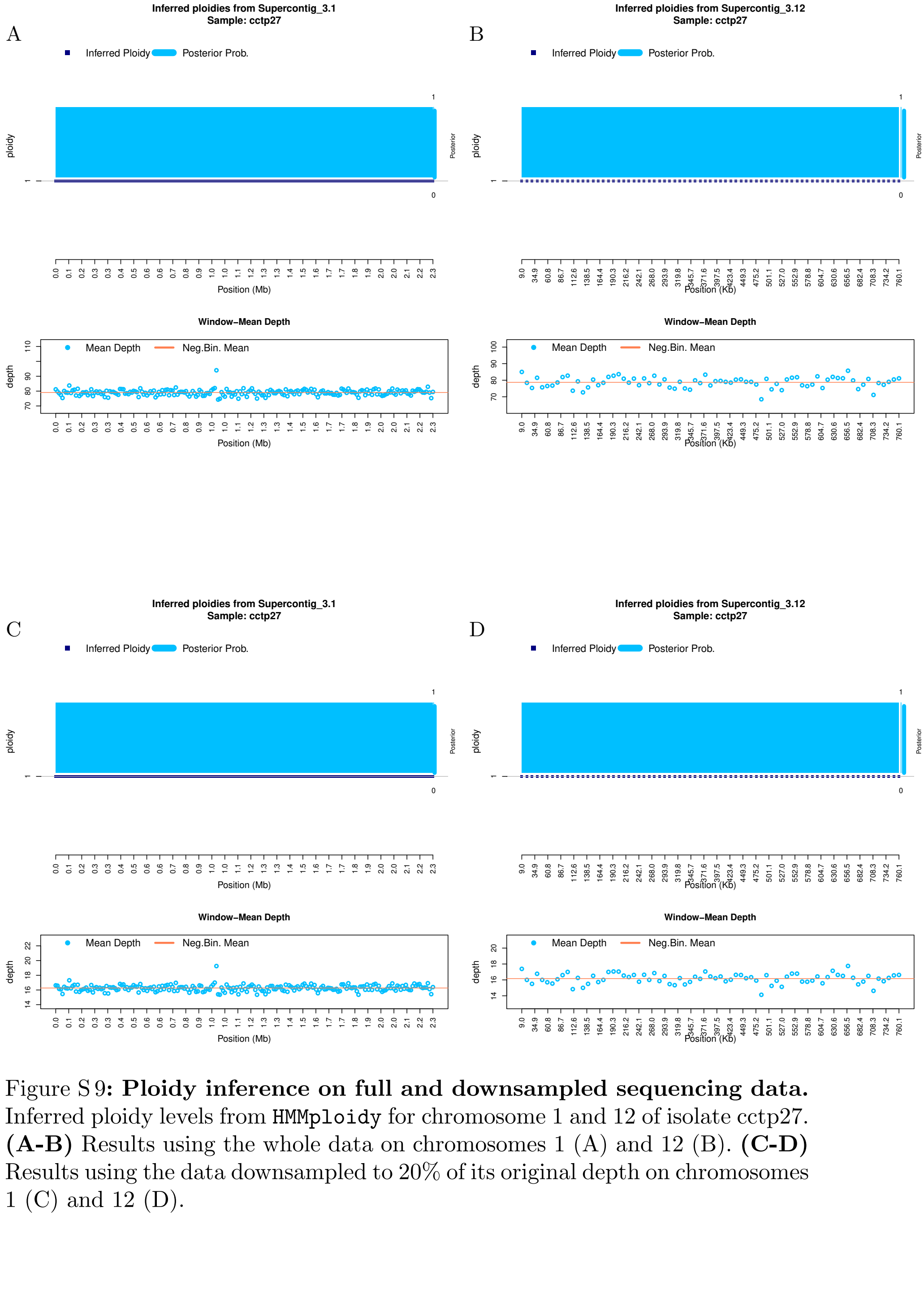


Figure S 9**: Ploidy inference on full and downsampled sequencing data.** Inferred ploidy levels from `HMMploidy` for chromosome 1 and 12 of isolate cctp27. **(A-B)** Results using the whole data on chromosomes 1 (A) and 12 (B). **(C-D)** Results using the data downsampled to 20% of its original depth on chromosomes 1 (C) and 12 (D).

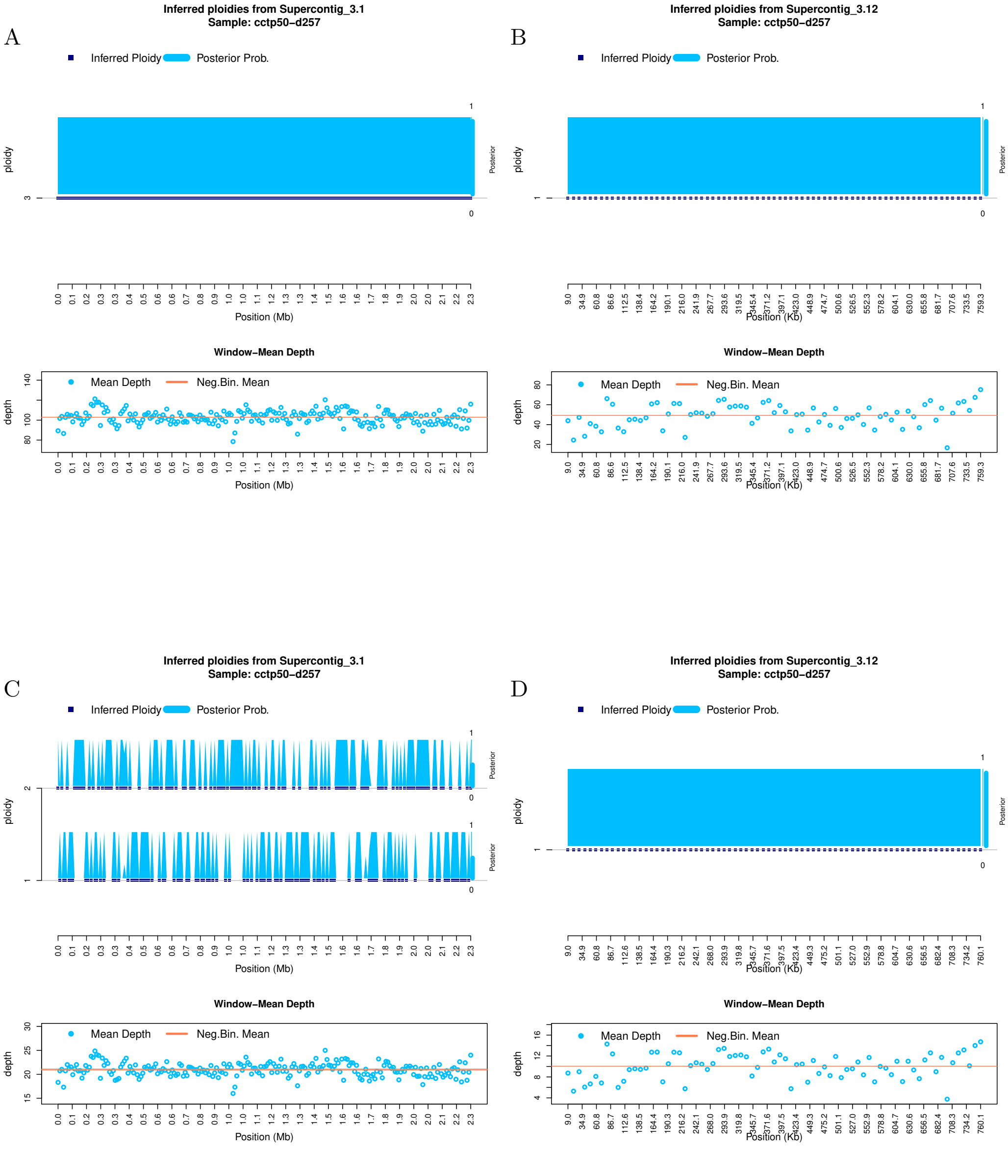


Figure S 10**: Ploidy inference on full and downsampled sequencing data.** Inferred ploidy levels from `HMMploidy` for chromosome 1 and 12 of isolate cctp50-d257. **(A-B)** Results using the whole data on chromosomes 1 (A) and 12 (B). **(C-D)** Results using the data downsampled to 20% of its original depth on chromosomes 1 (C) and 12 (D).

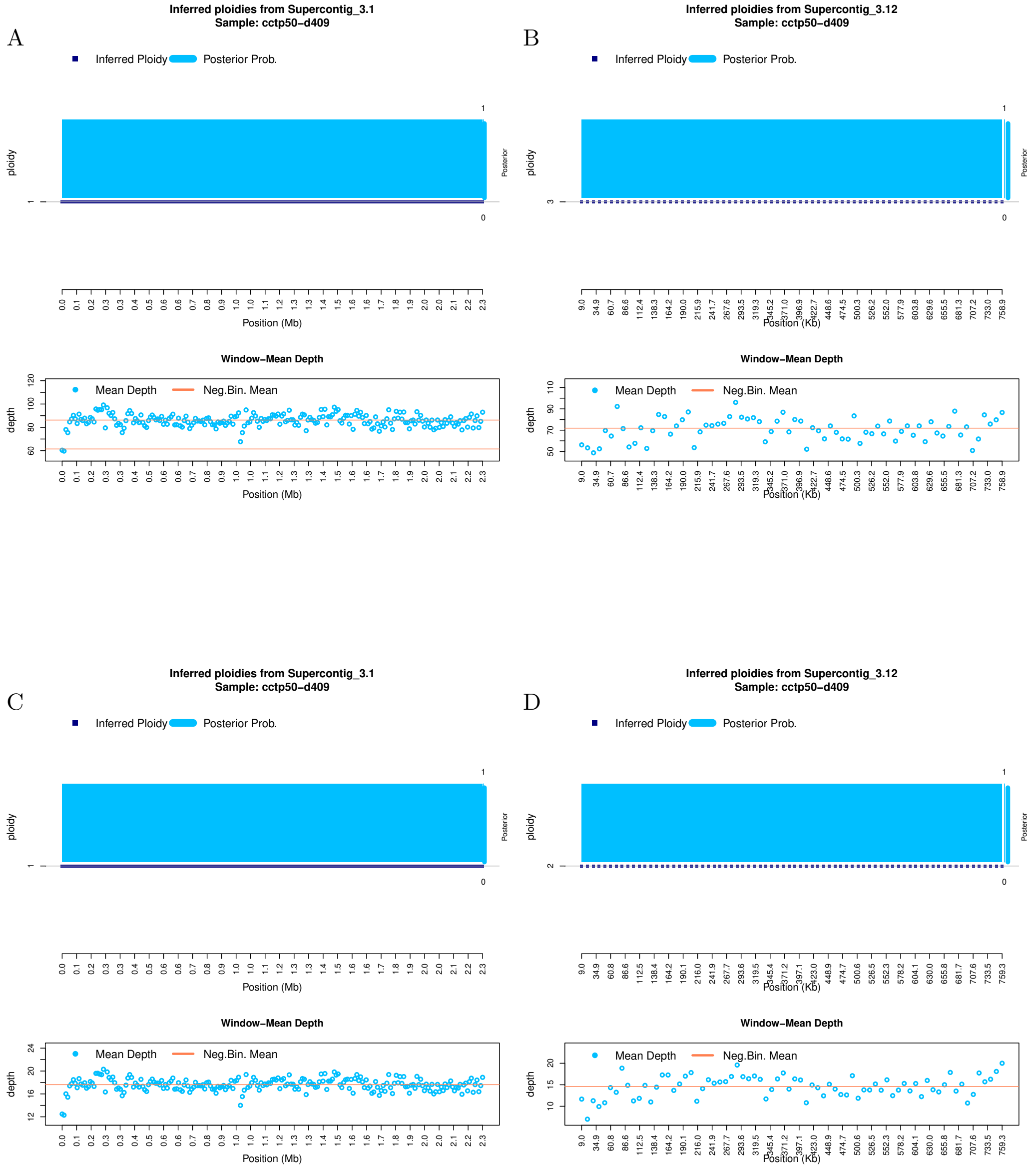


Figure S 11: **Ploidy inference on full and downsampled sequencing data.** Inferred ploidy levels from `HMMploidy` for chromosome 1 and 12 of isolate cctp50-d409. **(A-B)** Results using the whole data on chromosomes 1 (A) and 12 (B). **(C-D)** Results using the data downsampled to 20% of its original depth on chromosomes 1 (C) and 12 (D).

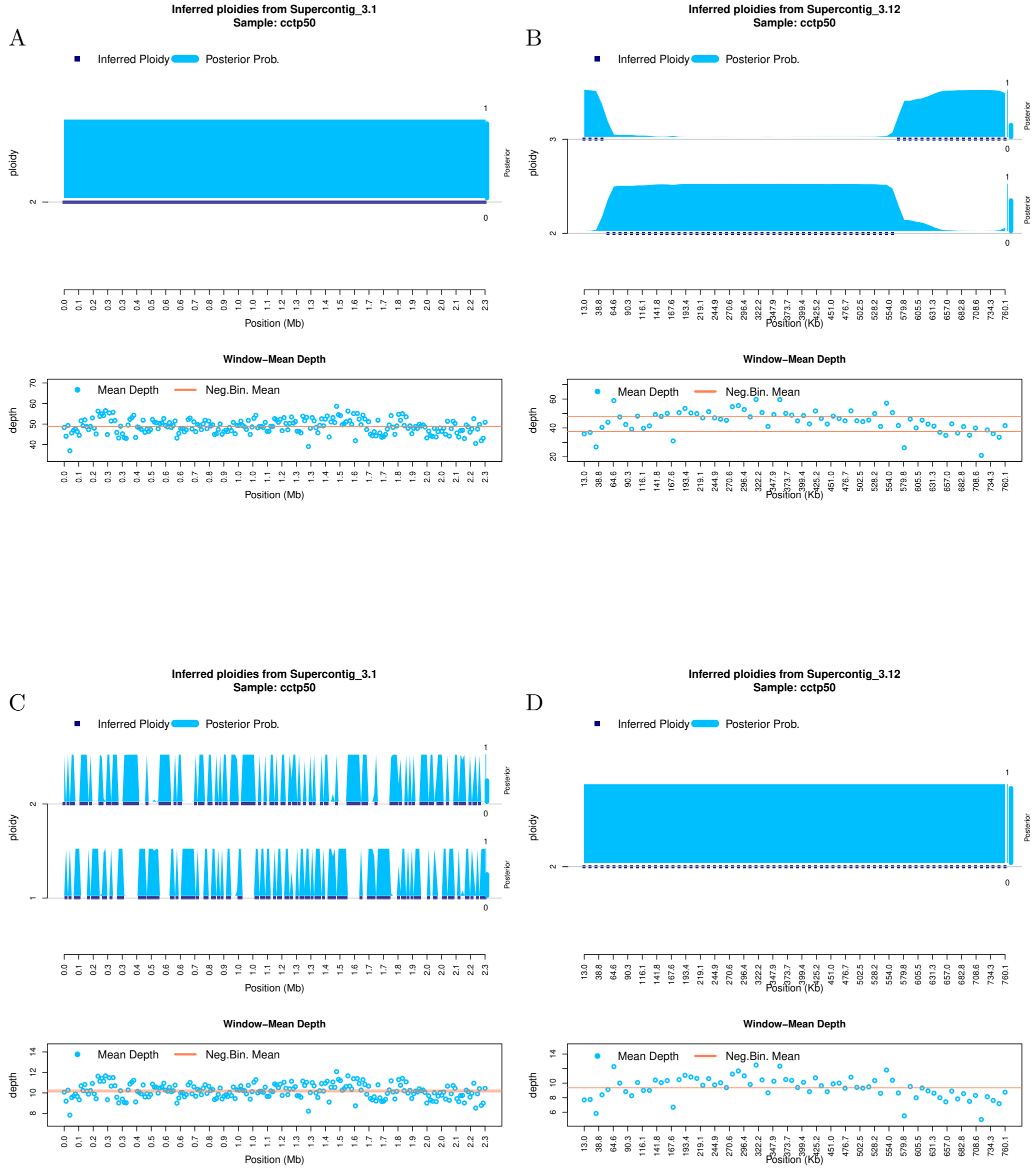


Figure S 12: **Ploidy inference on full and downsampled sequencing data.** Inferred ploidy levels from `HMMploidy` for chromosome 1 and 12 of isolate cctp50. **(A-B)** Results using the whole data on chromosomes 1 (A) and 12 (B). **(C-D)** Results using the data downsampled to 20% of its original depth on chromosomes 1 (C) and 12 (D).

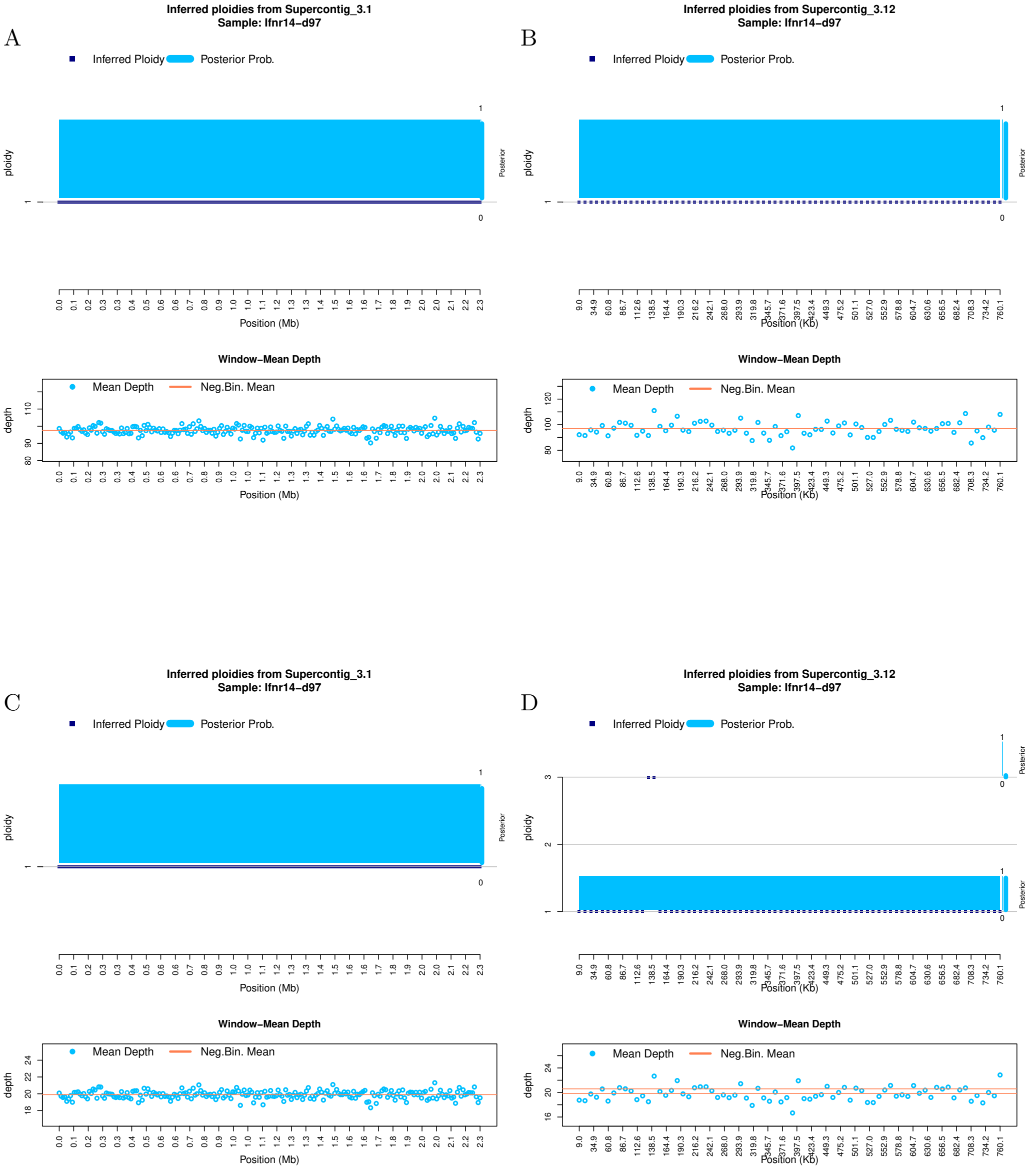


Figure S 13**: Ploidy inference on full and downsampled sequencing data.** Inferred ploidy levels from `HMMploidy` for chromosome 1 and 12 of isolate ifnr14-d97. **(A-B)** Results using the whole data on chromosomes 1 (A) and 12 (B). **(C-D)** Results using the data downsampled to 20% of its original depth on chromosomes 1 (C) and 12 (D).

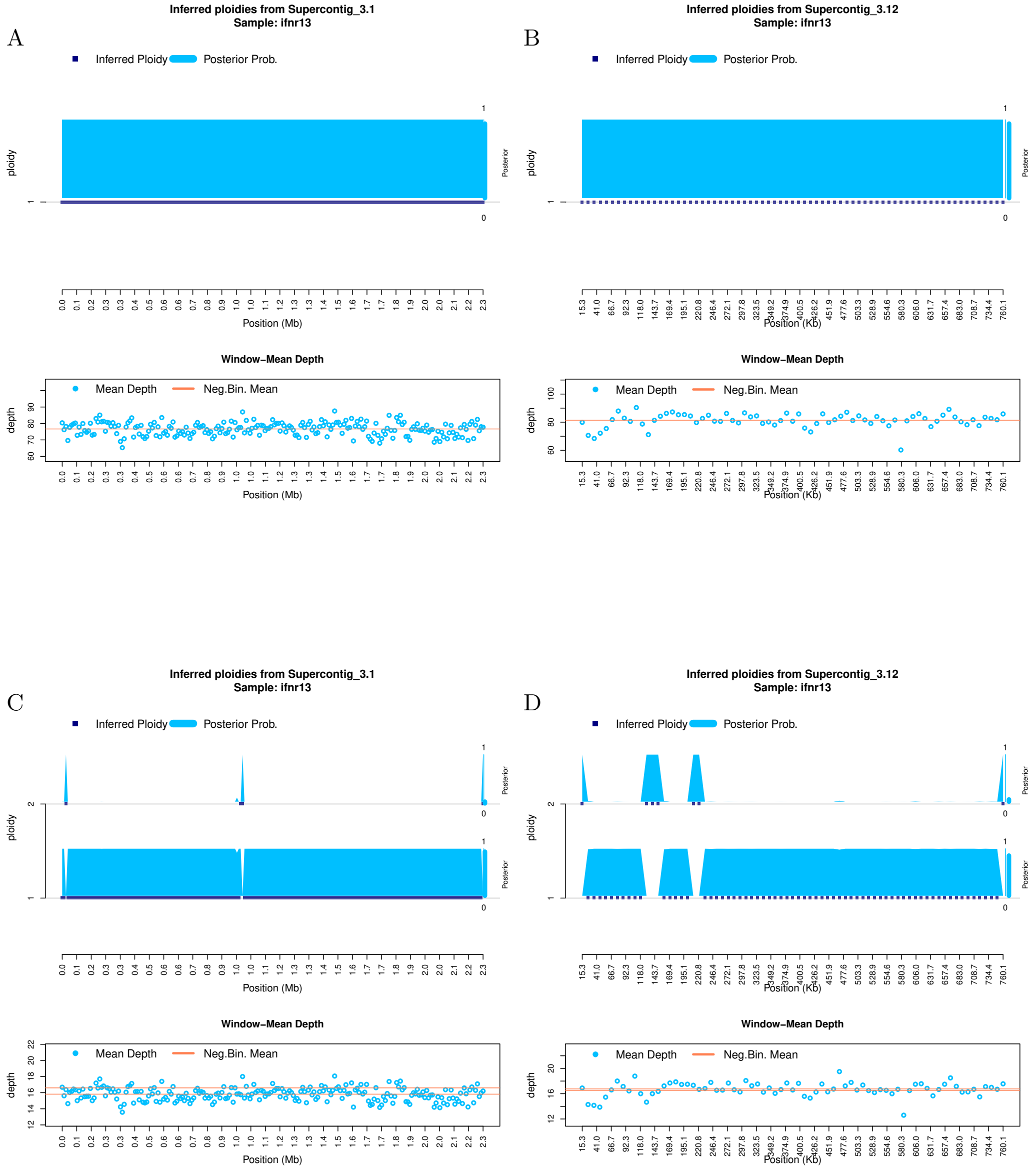


Figure S 14**: Ploidy inference on full and downsampled sequencing data.** Inferred ploidy levels from `HMMploidy` for chromosome 1 and 12 of isolate ifnr13. **(A-B)** Results using the whole data on chromosomes 1 (A) and 12 (B). **(C-D)** Results using the data downsampled to 20% of its original depth on chromosomes 1 (C) and 12 (D).

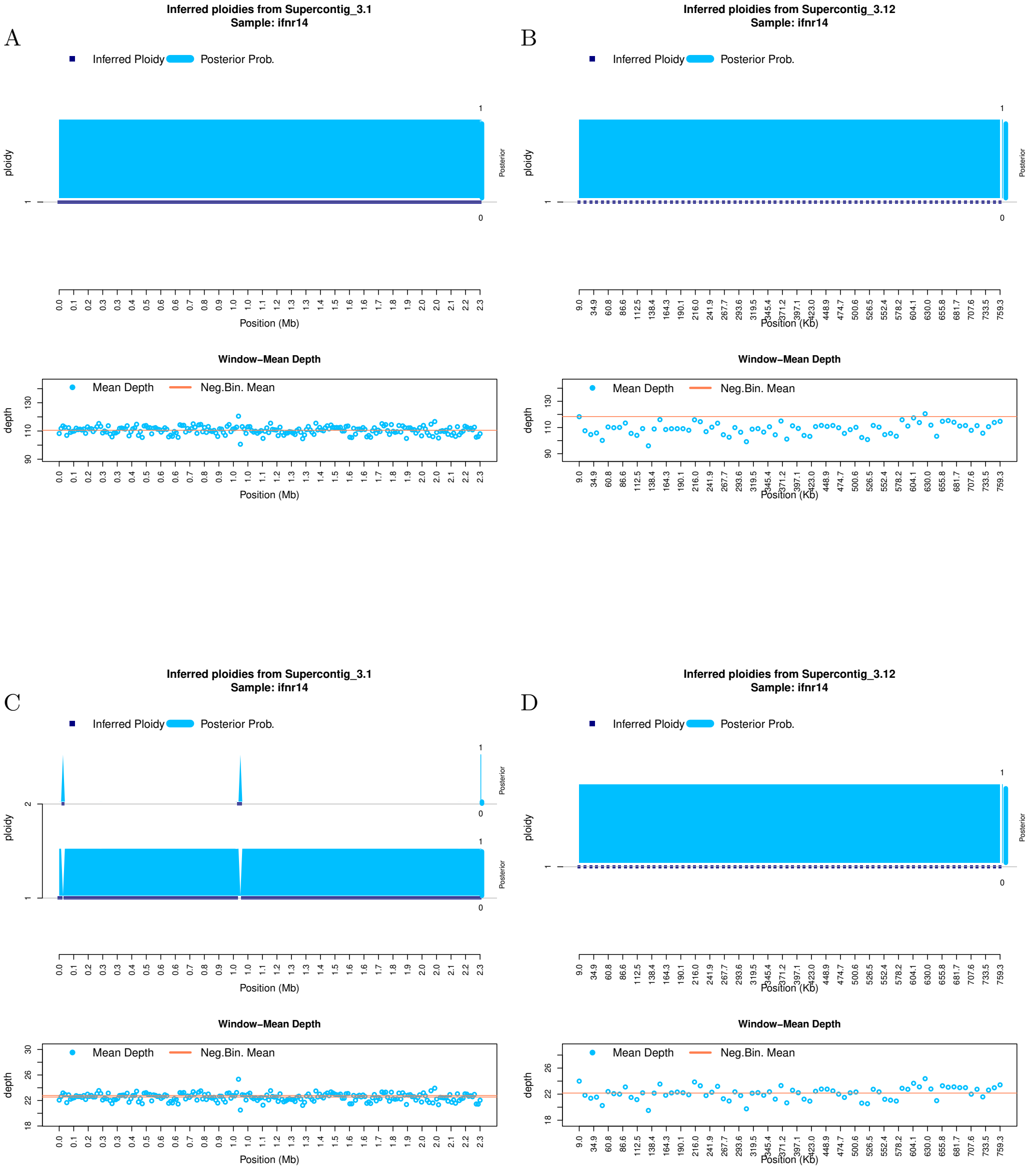


Figure S 15**: Ploidy inference on full and downsampled sequencing data.** Inferred ploidy levels from `HMMploidy` for chromosome 1 and 12 of isolate ifnr14. **(A-B)** Results using the whole data on chromosomes 1 (A) and 12 (B). **(C-D)** Results using the data downsampled to 20% of its original depth on chromosomes 1 (C) and 12 (D).

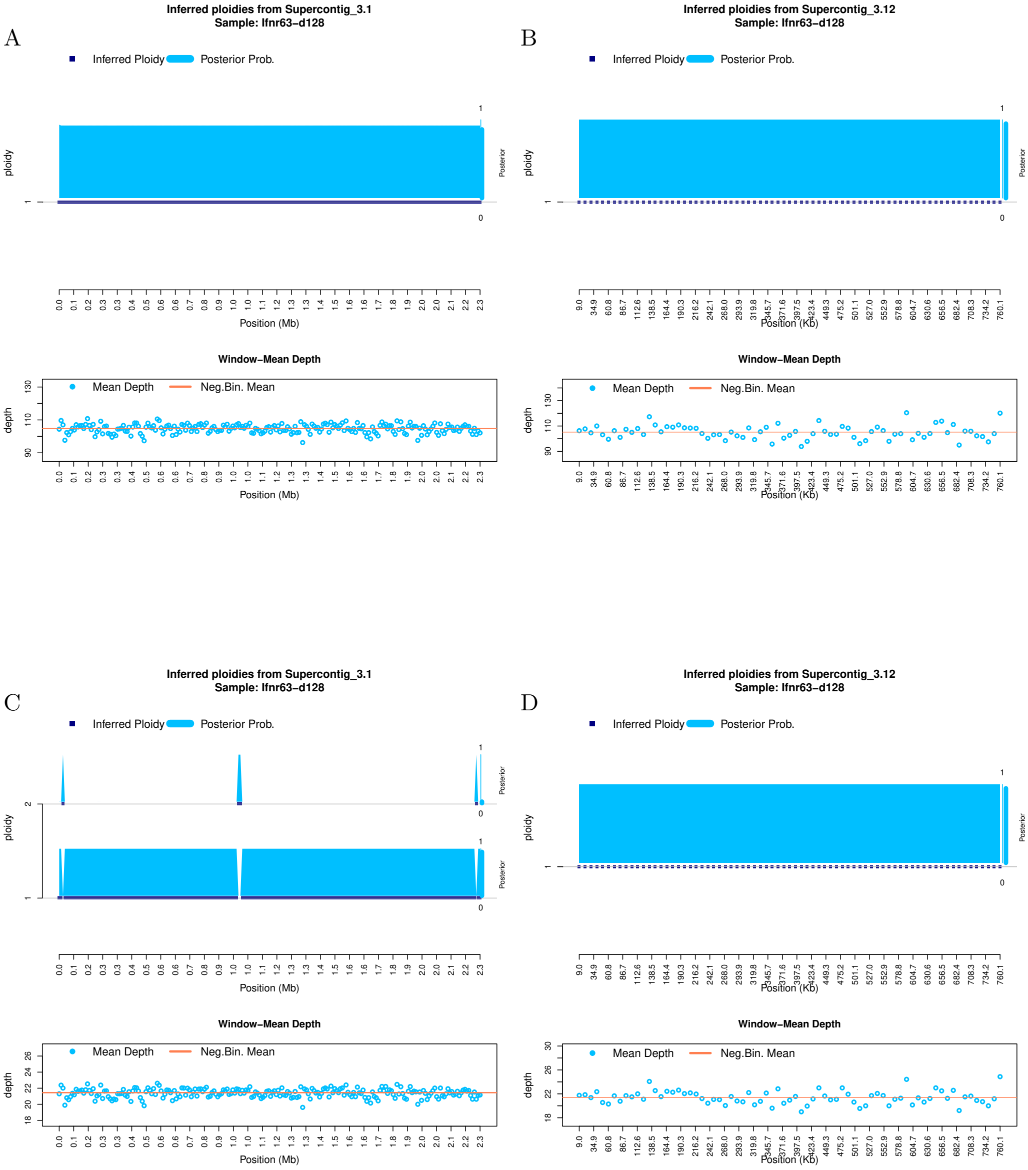


Figure S 16: **Ploidy inference on full and downsampled sequencing data.** Inferred ploidy levels from `HMMploidy` for chromosome 1 and 12 of isolate ifnr63-d128. **(A-B)** Results using the whole data on chromosomes 1 (A) and 12 (B). **(C-D)** Results using the data downsampled to 20% of its original depth on chromosomes 1 (C) and 12 (D).

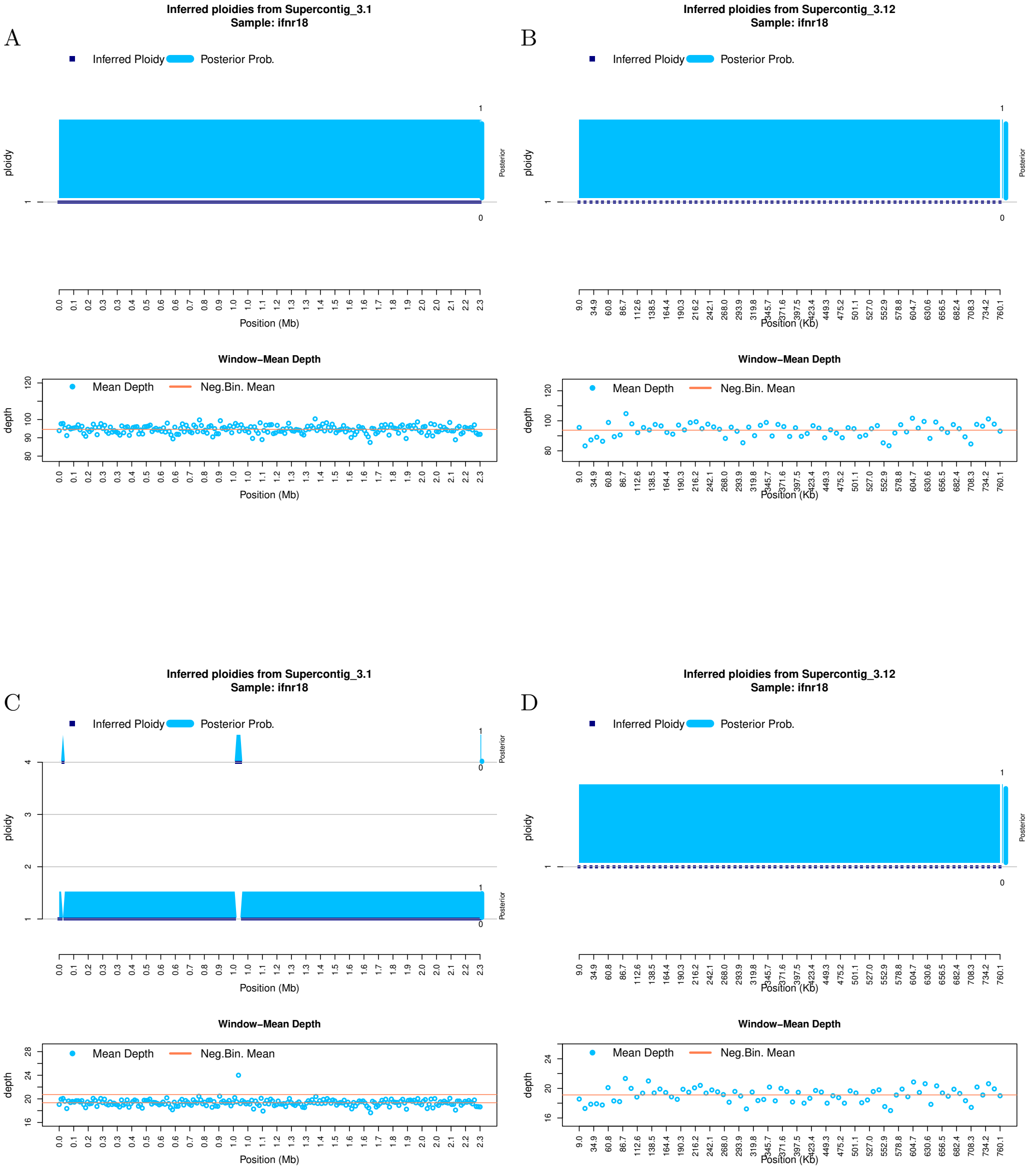


Figure S 17**: Ploidy inference on full and downsampled sequencing data.** Inferred ploidy levels from `HMMploidy` for chromosome 1 and 12 of isolate ifnr18. **(A-B)** Results using the whole data on chromosomes 1 (A) and 12 (B). **(C-D)** Results using the data downsampled to 20% of its original depth on chromosomes 1 (C) and 12 (D).

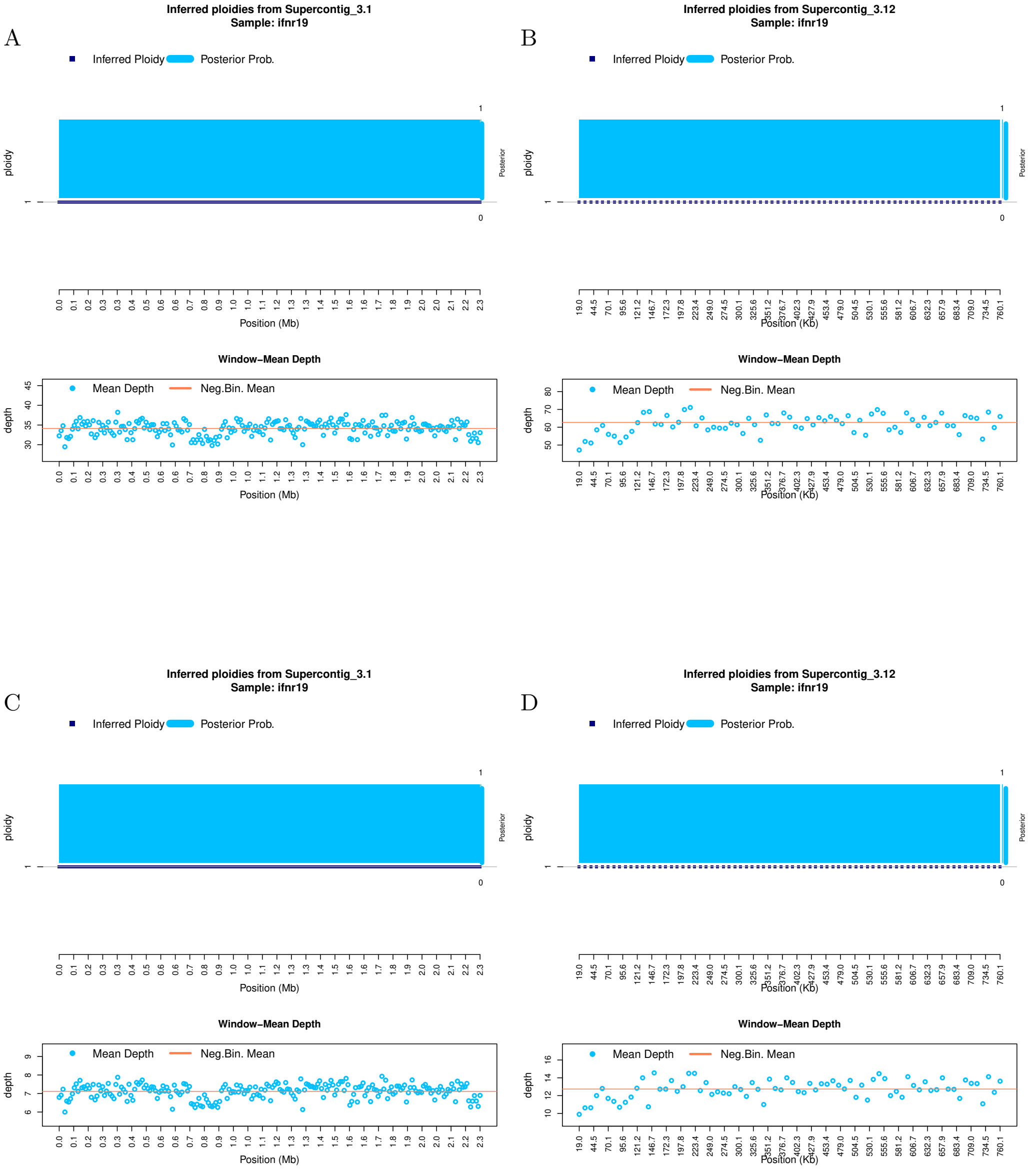


Figure S 18: **Ploidy inference on full and downsampled sequencing data.** Inferred ploidy levels from `HMMploidy` for chromosome 1 and 12 of isolate ifnr19. **(A-B)** Results using the whole data on chromosomes 1 (A) and 12 (B). **(C-D)** Results using the data downsampled to 20% of its original depth on chromosomes 1 (C) and 12 (D).

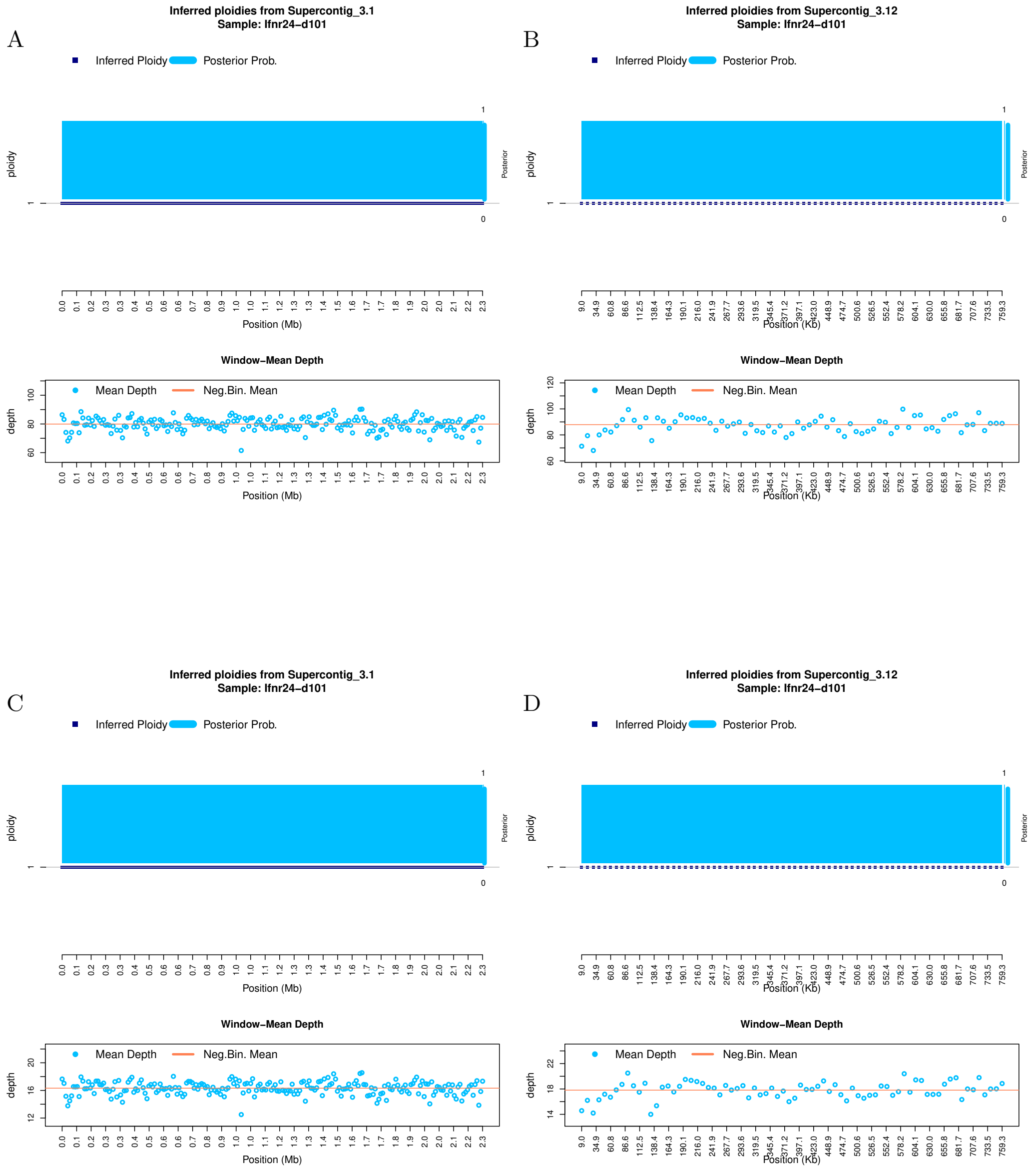


Figure S 19: **Ploidy inference on full and downsampled sequencing data.** Inferred ploidy levels from `HMMploidy` for chromosome 1 and 12 of isolate ifnr24-d101. **(A-B)** Results using the whole data on chromosomes 1 (A) and 12 (B). **(C-D)** Results using the data downsampled to 20% of its original depth on chromosomes 1 (C) and 12 (D).

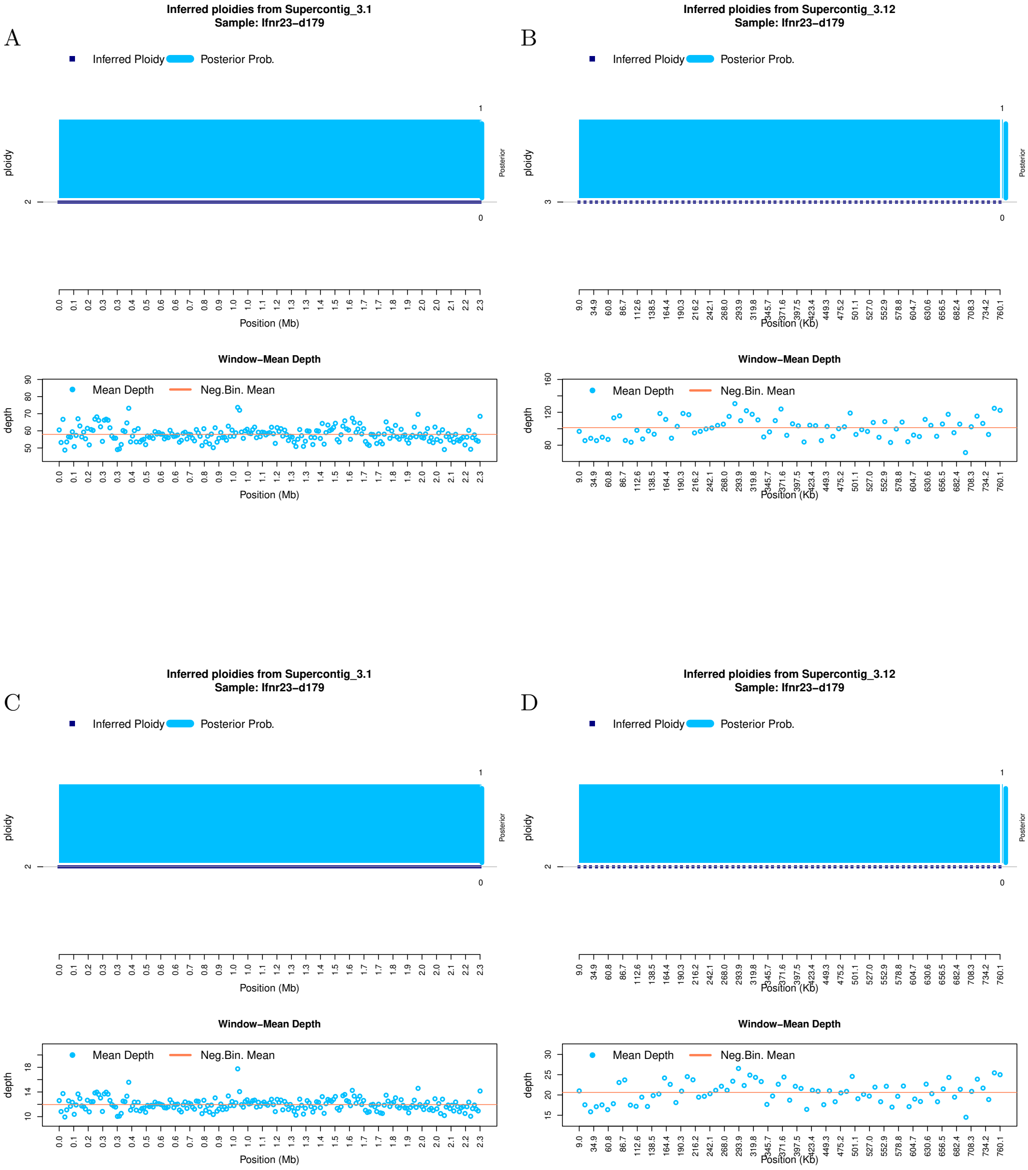


Figure S 20**: Ploidy inference on full and downsampled sequencing data.** Inferred ploidy levels from `HMMploidy` for chromosome 1 and 12 of isolate ifnr23-d179. **(A-B)** Results using the whole data on chromosomes 1 (A) and 12 (B). **(C-D)** Results using the data downsampled to 20% of its original depth on chromosomes 1 (C) and 12 (D).

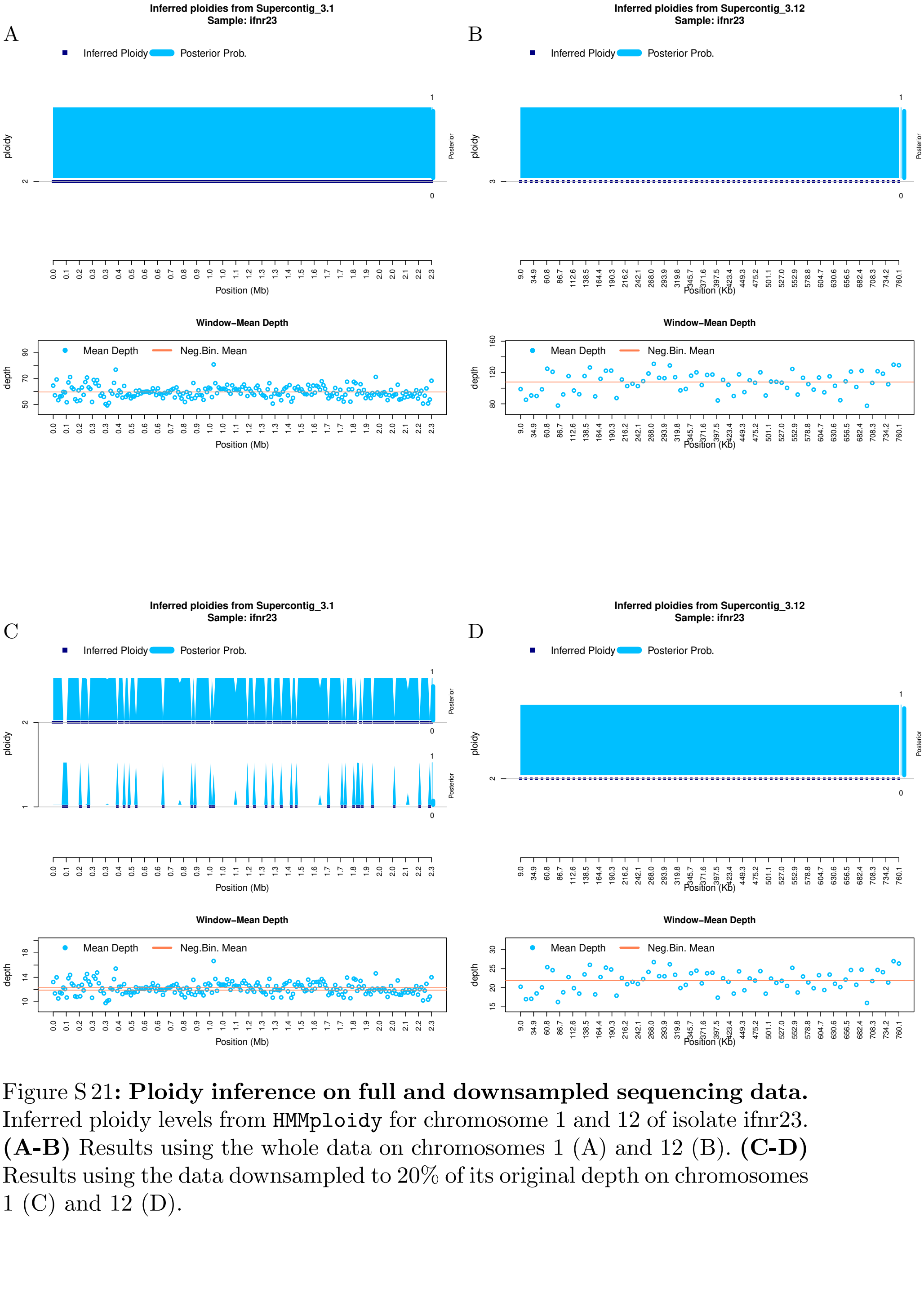


Figure S 21**: Ploidy inference on full and downsampled sequencing data.** Inferred ploidy levels from `HMMploidy` for chromosome 1 and 12 of isolate ifnr23. **(A-B)** Results using the whole data on chromosomes 1 (A) and 12 (B). **(C-D)** Results using the data downsampled to 20% of its original depth on chromosomes 1 (C) and 12 (D).

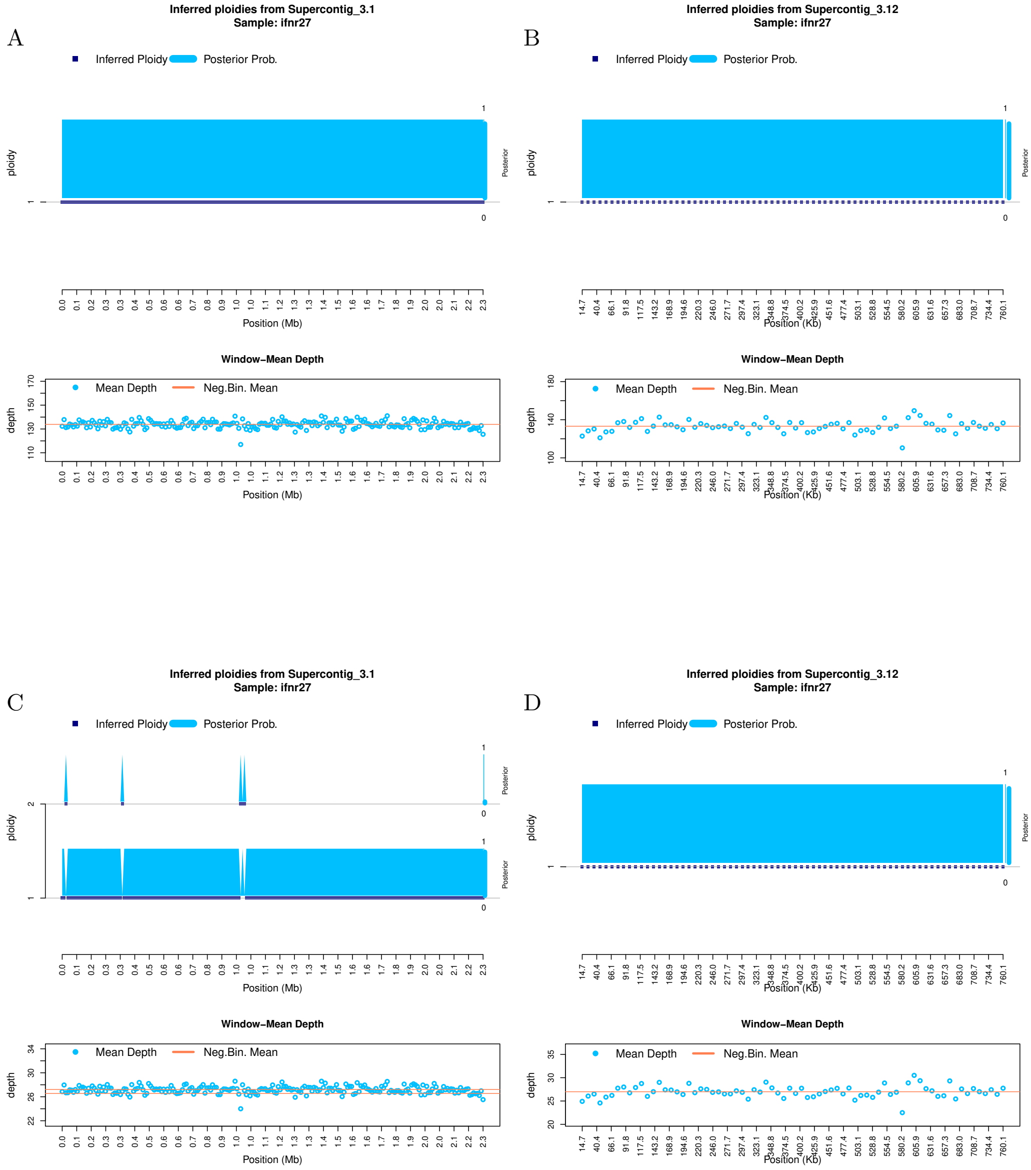


Figure S 22**: Ploidy inference on full and downsampled sequencing data.** Inferred ploidy levels from `HMMploidy` for chromosome 1 and 12 of isolate ifnr27. **(A-B)** Results using the whole data on chromosomes 1 (A) and 12 (B). **(C-D)** Results using the data downsampled to 20% of its original depth on chromosomes 1 (C) and 12 (D).

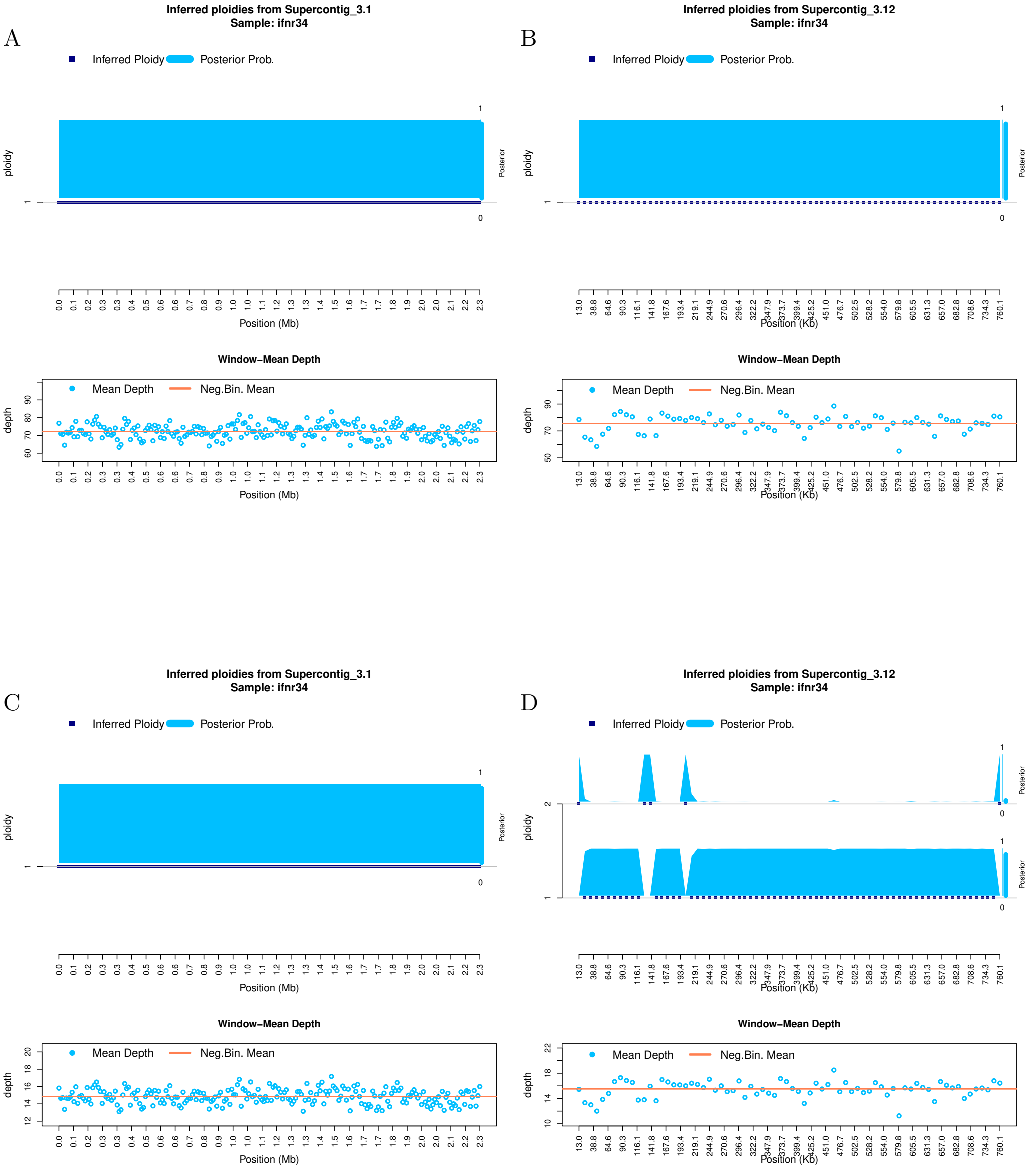


Figure S 23: **Ploidy inference on full and downsampled sequencing data.** Inferred ploidy levels from `HMMploidy` for chromosome 1 and 12 of isolate ifnr34. **(A-B)** Results using the whole data on chromosomes 1 (A) and 12 (B). **(C-D)** Results using the data downsampled to 20% of its original depth on chromosomes 1 (C) and 12 (D).

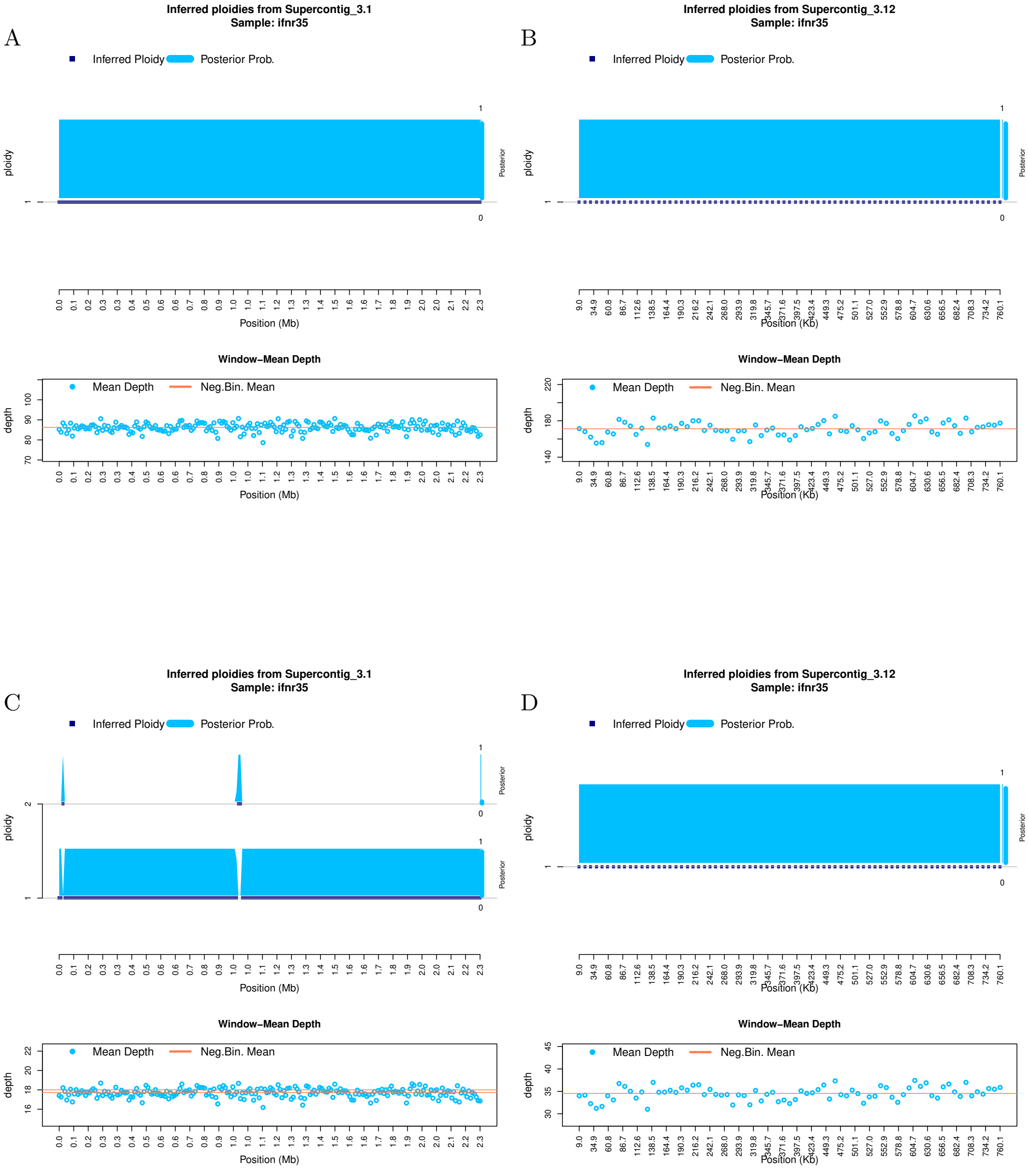


Figure S 24: **Ploidy inference on full and downsampled sequencing data.** Inferred ploidy levels from `HMMploidy` for chromosome 1 and 12 of isolate ifnr35. **(A-B)** Results using the whole data on chromosomes 1 (A) and 12 (B). **(C-D)** Results using the data downsampled to 20% of its original depth on chromosomes 1 (C) and 12 (D).

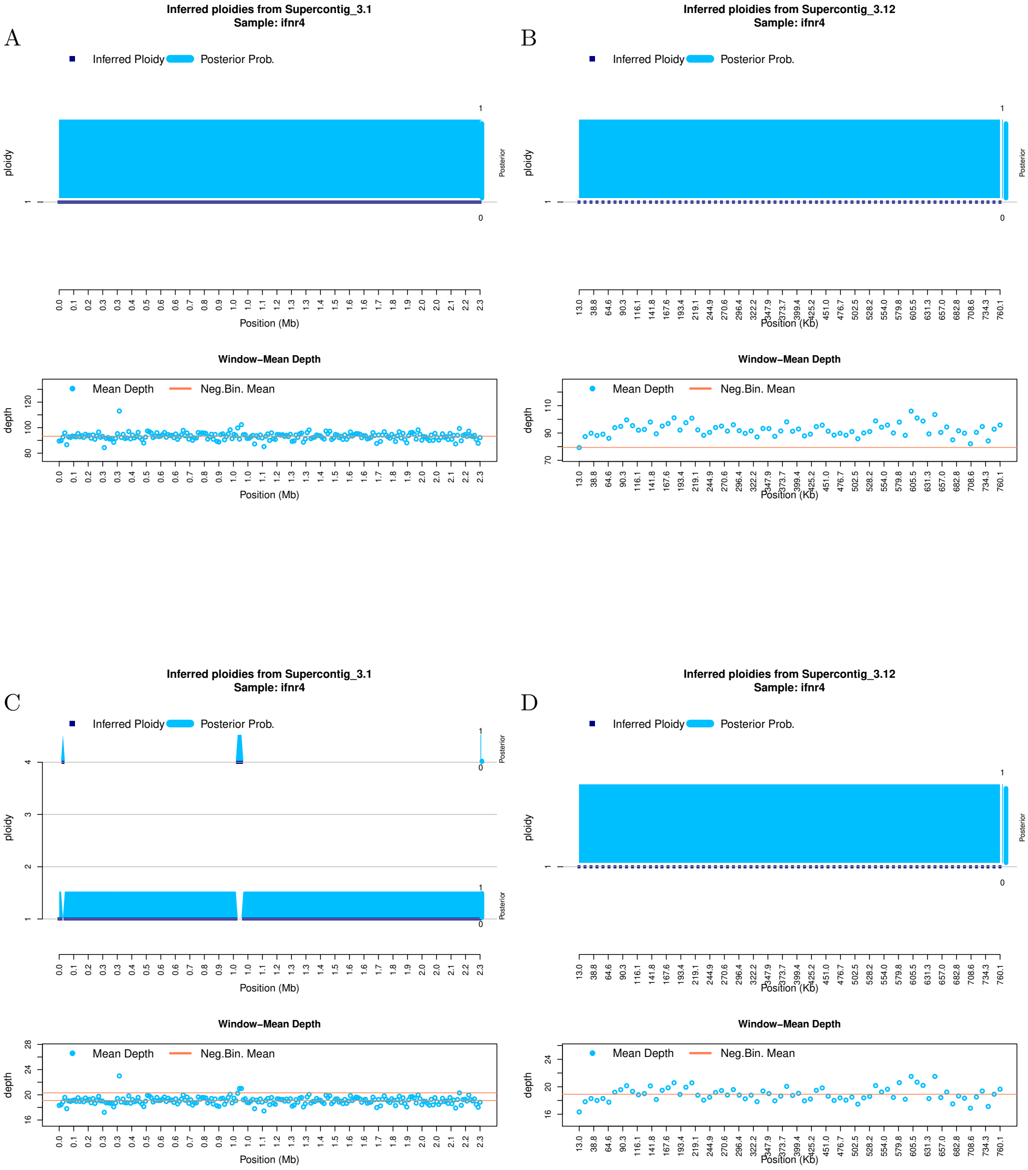


Figure S 25**: Ploidy inference on full and downsampled sequencing data.** Inferred ploidy levels from `HMMploidy` for chromosome 1 and 12 of isolate ifnr4. **(A-B)** Results using the whole data on chromosomes 1 (A) and 12 (B). **(C-D)** Results using the data downsampled to 20% of its original depth on chromosomes 1 (C) and 12 (D).

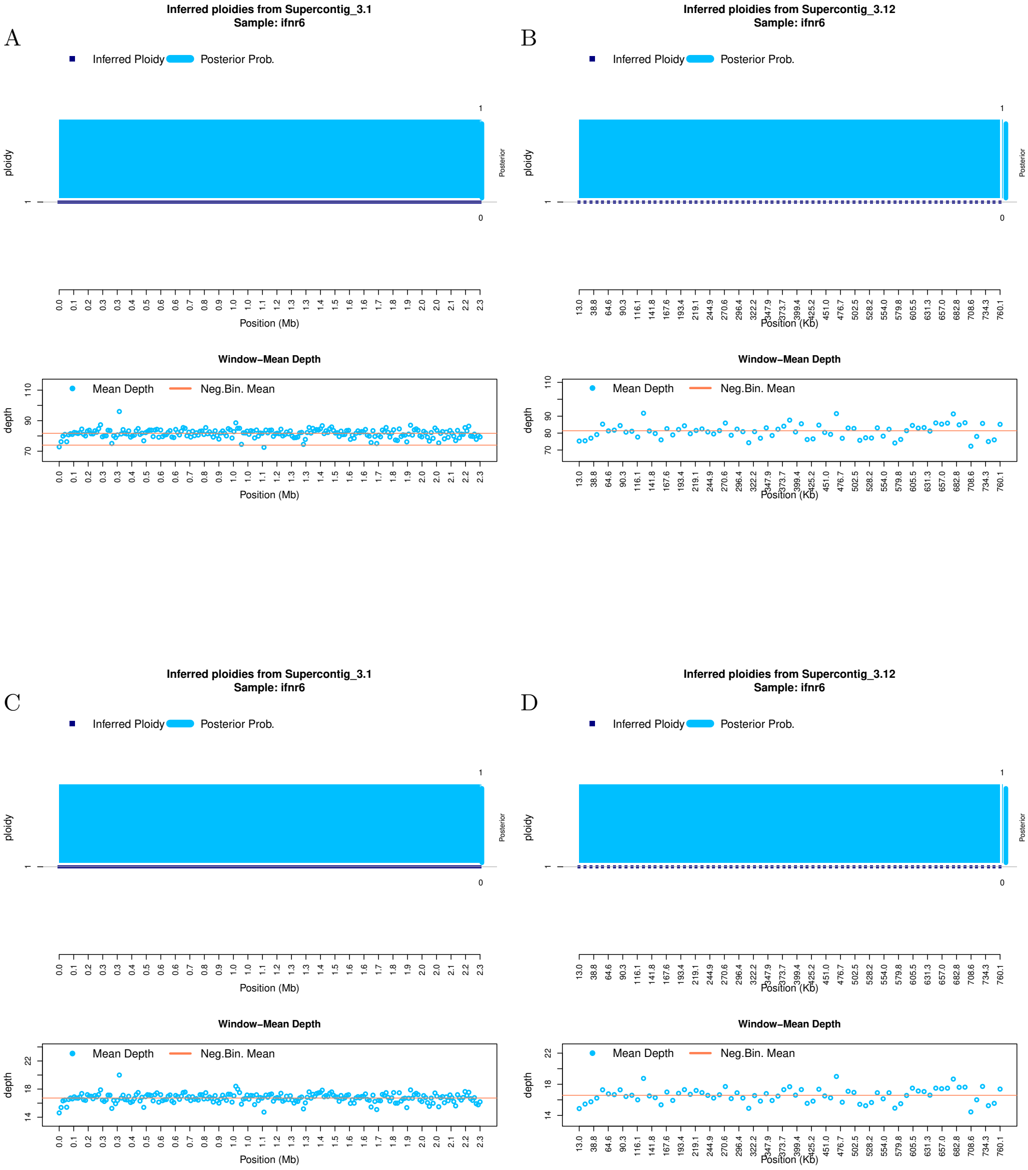


Figure S 26**: Ploidy inference on full and downsampled sequencing data.** Inferred ploidy levels from `HMMploidy` for chromosome 1 and 12 of isolate ifnr6. **(A-B)** Results using the whole data on chromosomes 1 (A) and 12 (B). **(C-D)** Results using the data downsampled to 20% of its original depth on chromosomes 1 (C) and 12 (D).

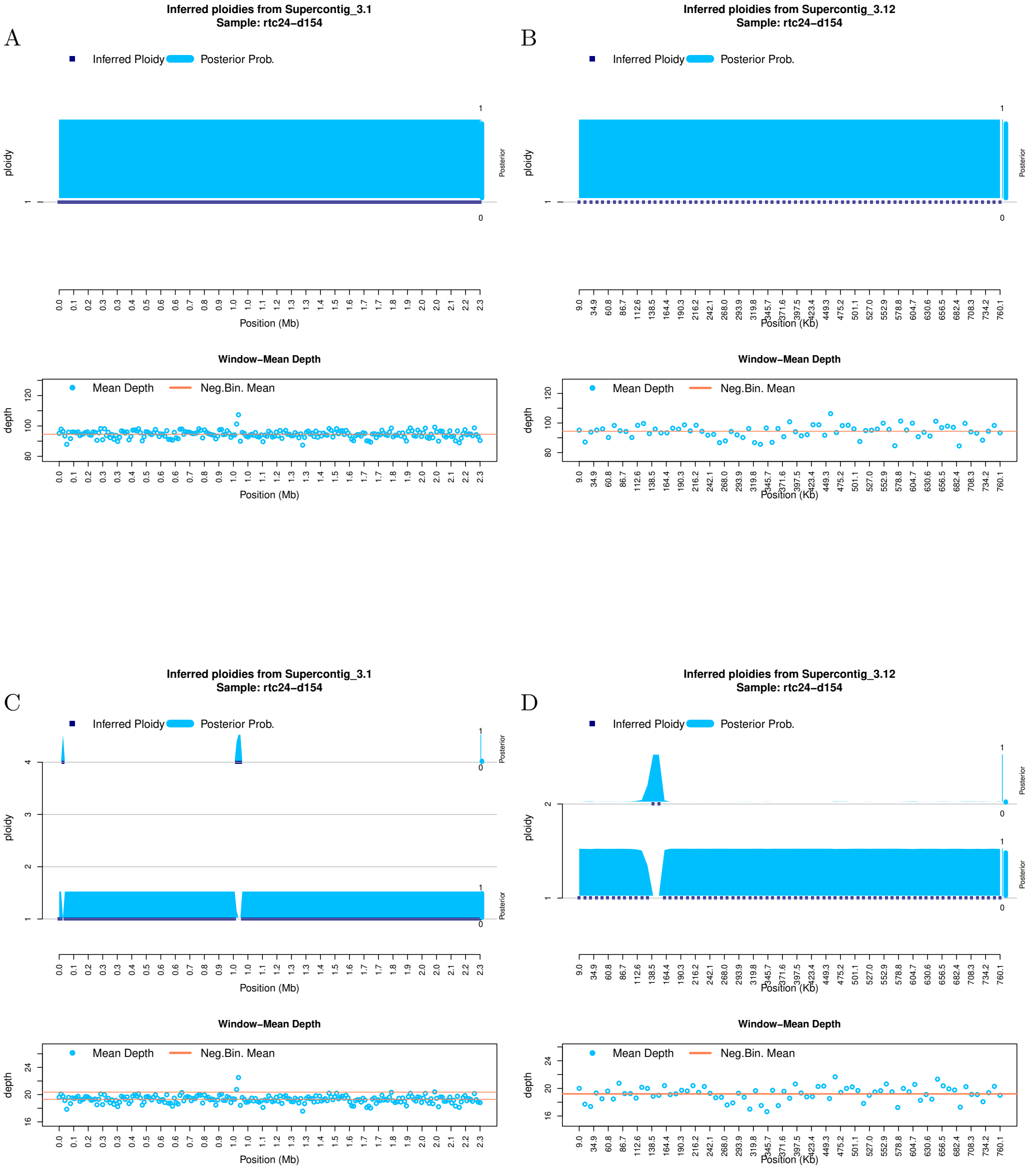


Figure S 27: **Ploidy inference on full and downsampled sequencing data.** Inferred ploidy levels from `HMMploidy` for chromosome 1 and 12 of isolate rtc24-d154. **(A-B)** Results using the whole data on chromosomes 1 (A) and 12 (B). **(C-D)** Results using the data downsampled to 20% of its original depth on chromosomes 1 (C) and 12 (D).

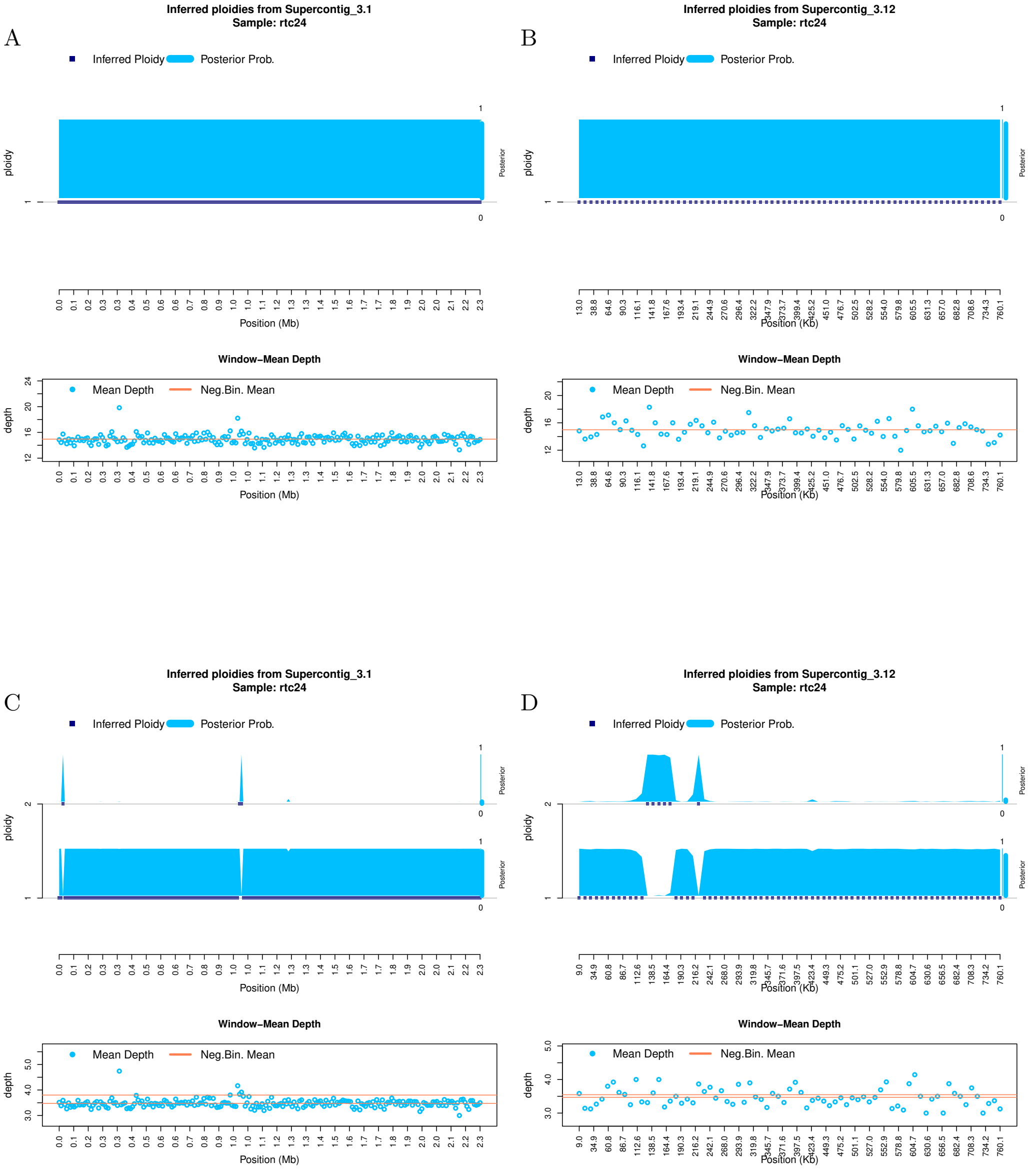


Figure S 28: **Ploidy inference on full and downsampled sequencing data.** Inferred ploidy levels from `HMMploidy` for chromosome 1 and 12 of isolate rtc24. **(A-B)** Results using the whole data on chromosomes 1 (A) and 12 (B). **(C-D)** Results using the data downsampled to 20% of its original depth on chromosomes 1 (C) and 12 (D).

[42] Rhodes, J., Beale, M.A., Vanhove, M., Jarvis, J.N., Kannambath, S., Simpson, J.A., Ryan, A., Meintjes, G., Harrison, T.S., Fisher, M.C., Bicanic, T.: A Population Genomics Approach to Assessing the Genetic Basis of Within-Host Microevolution Underlying Recurrent Cryptococcal Meningitis Infection. G3 Genes—Genomes—Genetics

(2017). https://doi.org/10.1534/g3.116.037499, https://doi.org/10.1534/g3.116.037499

[43] Rhodes, J., Desjardins, C.A., Sykes, S.M., Beale, M.A., Vanhove, M., Sakthikumar, S., Chen, Y., Gujja, S., Saif, S., Chowdhary, A., Lawson, D.J., Ponzio, V., Colombo, A.L., Meyer, W., Engelthaler, D.M., Hagen, F., Illnait-Zaragozi, M.T., Alanio, A., Vreulink, J.M., Heitman, J., Perfect, J.R., Litvintseva, A.P., Bicanic, T., Harrison, T.S., Fisher, M.C., Cuomo, C.A.: Tracing Genetic Exchange and Biogeography of Cryptococcus neoformans var. grubii at the Global Population Level. Genetics **207**(1), 327–346 (07 2017). https://doi.org/10.1534/genetics.117.203836, https://doi.org/10.1534/genetics.117.203836
